# Chronic high dietary sucrose induces sexually dimorphic metabolic adaptations in liver and adipose tissue

**DOI:** 10.1101/2020.05.20.106922

**Authors:** Erin J Stephenson, Amanda S Stayton, Aarti Sethuraman, Prahlad K Rao, Charles Klazer Gomes, Molly C Mulcahy, Liam McAllan, Michelle A Puchowicz, Joseph F Pierre, Dave Bridges, Joan C Han

**Author notes:** Corresponding author. **Contact Info**, Correspondence.

## Abstract

Almost all effective treatments for non-alcoholic fatty liver disease (NAFLD) involve reduction of adiposity, which suggests the metabolic axis between liver and adipose tissue is essential to NAFLD development. Since excessive dietary sugar intake may be an initiating factor for NAFLD, we have characterized the metabolic effects of liquid sucrose intake at concentrations relevant to typical human consumption in mice. We report that sucrose intake induces sexually dimorphic effects in liver, adipose tissue, and the microbiome; differences concordant with steatosis severity. We show that when steatosis is decoupled from impairments in insulin responsiveness, sex is a moderating factor that influences sucrose-driven lipid storage and the contribution of *de novo* fatty acid synthesis to the overall hepatic triglyceride pool. Our findings provide physiologic insight into how sex influences the regulation of adipose-liver crosstalk and highlight the importance of extrahepatic metabolism in the pathogenesis of diet-induced steatosis and NAFLD.

## Introduction

Non-alcoholic fatty liver disease (NAFLD) is the most common form of chronic liver disease, affecting ~25% of Earth’s human population (Younossi et al., 2016). NAFLD is characterized by the presence of >5% steatosis (simple steatosis), with or without inflammation and/or scarring in the liver (nonalcoholic steatohepatitis; NASH) (Sanyal, 2019). Simple steatosis progresses to NASH in 59% of patients (Younossi et al., 2016), whereas advancing fibrosis in NASH can lead to cirrhosis, hepatocellular carcinoma and liver failure (Sanyal, 2019). Approximately 50% of all NAFLD patients have obesity (Younossi et al., 2016), and individuals with NAFLD have a higher incidence of type 2 diabetes, cardiovascular disease and all-cause mortality (Armstrong et al., 2014; Kim et al., 2013; Younossi et al., 2016). However, not all patients with NAFLD have other metabolic comorbidities (and vice-versa), suggesting that although NAFLD and other metabolic disease states are related, their etiology can be decoupled (Shi et al., 2020).

Findings from observational studies and meta analyses indicate that excessive sugar intake or compromised sugar metabolism may be an initiating factor for NAFLD in humans (Arenaza et al., 2019; Asgari-Taee et al., 2019; Assy et al., 2008; Cahlin et al., 1973; Chen et al., 2019; Ma et al., 2015). Although dietary sugars have been used extensively to induce cardiometabolic abnormalities in animal models (Bacon et al., 1984; Bukowiecki et al., 1983; Burke et al., 2018; Chen et al., 2019; Kawasaki et al., 2009; Oliveira et al., 2014; Ritze et al., 2014; Softic et al., 2019; Togo et al., 2019), the molecular mechanisms underlying sugar-induced hepatic lipid accumulation remain poorly understood in a context that is translational to human dietary practices (Choo and Sievenpiper, 2015). Most preclinical studies rely upon use of supraphysiologic concentrations of sucrose, glucose or fructose to induce steatosis and other metabolic comorbidities (Chen et al., 2011; Choo and Sievenpiper, 2015; Softic et al., 2019; Soria et al., 2001), often with the confounding addition of excess dietary lipid (Chen et al., 2011; Oliveira et al., 2014; Softic et al., 2019). This lack of translatability, in combination with reports that the effects of dietary sugars are dependent on physical form (i.e., liquid versus solid) (Ritze et al., 2014; Togo et al., 2019), make it difficult to draw mechanistic conclusions applicable to how sugar intake leads to NAFLD in humans.

Nonetheless, it has been generally accepted that high sucrose consumption (or consumption of the not-meaningfully different high-fructose corn syrup) causes an elevation in *de novo* fatty acid synthesis (and, subsequently, hepatic triglyceride storage) due to the ability of fructose, one half of the sucrose disaccharide, to proceed directly toward *de novo* fatty acid synthesis (Mayes, 1993; Spruss and Bergheim, 2009). However, tracer studies at physiologic fructose concentrations indicate that only a small percentage (<1%) of ingested fructose is directly converted to lipid (most becomes glucose or lactate) (Laughlin, 2014; Sun and Empie, 2012). Thus, in addition to fructose-driven fatty acid synthesis in liver, the development of steatosis following high sucrose intake likely involves utilization of fatty acids derived from extrahepatic sources (Chong et al., 2007; Hellerstein, 1999; Parks et al., 2008).

Most circulating non-esterified fatty acids (NEFA) are derived from adipose tissue lipolysis (van Hall et al., 2002). Adipose tissue mass increases rapidly in response to sucrose intake (Bukowiecki et al., 1983; Chen et al., 2011; Oliveira et al., 2014), whereas increased adiposity following sucrose feeding in humans (Smith et al., 1973) and rats (Soria et al., 2001) leads to an increase in the rate of adipose tissue lipolysis *ex vivo.* Thus, in addition to direct effects on liver, high dietary sucrose intake induces molecular changes in adipose tissue that lead to dysregulated lipolysis and a subsequent increase circulating NEFA which would, in turn, be expected to increase demand for fatty acid uptake and re-esterification in the liver.

Here, we investigated the effects of chronic, physiologically-relevant dietary sucrose intake on energy balance and the liver-adipose axis. We hypothesized that chronic sucrose intake would drive hepatic steatosis indirectly, primarily as a result of dysregulated adipose tissue lipolysis and increased reesterification of extrahepatic derived fatty acids in the liver. Additionally, because there is a lack of *in vivo* data to support epidemiological data (Ballestri et al., 2017; Group. et al., 2015) and *in silico* modelling (Cvitanovic Tomas et al., 2018; Lonardo et al., 2019) indicating that NAFLD develops through distinct metabolic processes in males and females, we also sought to determine how sex impacts the severity of sucrose-induced steatosis and other metabolic parameters that may be affected by chronic liquid sucrose intake.

## Methods

### Generation of Mice and Experimental Design

Female mice homozygous for a LoxP-modified *Pnpla2* allele (B6N.129S-*Pnpla2*^tm1Eek/J^, stock number 024278 (Sitnick et al., 2013); *Pnpla2*^Fl/Fl^) and male mice hemizygous for an adipose tissue-specific Cre recombinase, under control of the adiponectin promotor (B6FVB-Tg^(*Adipoq*-cre)1Evdr/J^, stock number 028020 (Eguchi et al., 2011); Cre/^+^) mice were purchased from The Jackson Laboratory and crossed to generate breeding stock that were heterozygous for the targeted mutation of *Pnpla2*, with or without hemizygosity for the *Adipoq*-cre transgene (*Pnpla2*^Fl/+;^Cre/^+^ or *Pnpla2*^Fl/+;+/+^). The originating mice in the floxed strain were congenic to a C57BL/6NJ background, whereas the mice in the Cre strain were congenic to a C57BL6/J background. To generate experimental mice with or without adipose-specific deletion of exons 2-7 of *Pnpla2*, *Pnpla2*^Fl/+^;Cre/^+^ mice were mated with *Pnpla2*^Fl/+^;^+/+^ mice. Four of the resulting six genotypes were utilized in this study; the wild-type mice, with no genetic alterations (+/+;+/+), mice with adipose-specific deletion of *Pnpla2*, (*Pnpla2*^Fl/Fl^;Cre/^+^), and two additional control groups (*Pnpla2*^Fl/Fl;+/+^ and +/+;Cre/^+^). All mice were of a single generation and were genotyped at 14 days of age by PCR of tail DNA. Following genotyping, mice were randomly assigned to receive either water (control groups) or sucrose (10% w/v, prepared in distilled water) in their drinking water once they reached experimental age. Both male and female mice were weaned onto standard rodent chow (Envigo #7912) and water at 21-24 days of age, with their assigned experimental treatment beginning at 10 weeks of age. Mice were housed in ventilated cages with corn cobb bedding in a humidity and temperature-controlled environment at ~22°C with free access to food and water (or water containing 10% w/v sucrose). This concentration of sucrose was chosen because it closely approximates the concentration of sugar found across many readily available sugar-sweetened beverages made for human consumption (Ventura et al., 2011). The bulk of experiments were performed on mice of the +/+;+/+ genotype, in which the presence of the *Nnt* gene was verified by quantification of transcript levels in adipose tissue. In instances where *Pnpla2*^Fl/Fl^;Cre/^+^ mice were studied alongside all three control groups, the control group data (when not different between groups) were combined into a single group and labelled CON. Due to increased fluid intake of mice receiving sucrose, all cages were changed twice weekly. All procedures were performed in accordance with the National Institute of Health Guidelines for the Care and Use of Experimental animals and were pre-approved by the University of Tennessee Health Science Center Institutional Animal Care and Use Committee.

### Longitudinal Body Composition Monitoring

Mice were weighed and body composition was non-invasively measured using an EchoMRI 1100 (EchoMRI), beginning at 4 weeks of age, and was measured every week thereafter for the duration of the study.

### Indirect Calorimetry, Physical Activity Monitoring and Measurement of Food and Water intake

At 20 weeks of age, mice were individually housed in a home cage-style Comprehensive Laboratory Animal Monitoring System (CLAMS, Columbus Instruments), for two weeks (one week at 25°C, one week at 28°C). Following a 24hr acclimation period, VO_2_ and VCO_2_ were measured via open-circuit indirect calorimetry using Oxymax software (Columbus Instruments). Energy expenditure was calculated using the Lusk equation(Lusk, 1924). Total activity was calculated as the combined number of infrared beam breaks along the X-and Y-axes, whereas ambulatory activity was calculated as the combined number of consecutive X- and Y-axes beam breaks occurring in a single series. Food intake from suspended feeder baskets was determined via a linked load cell. Water/sucrose intake was measured manually, by regular weighing of the water bottles. Rates of fat and carbohydrate oxidation were calculated from indirect calorimetry data as previously described, assuming negligible protein oxidation (Peronnet and Massicotte, 1991). Both body fat and lean mass were included as covariates in the models used to analyze data obtained from the CLAMS experiments.

### Glucose and Insulin Tolerance Tests

Glucose and insulin tolerance were assessed at 24 weeks of age, with at least 4 days between tests. Mice were fasted for 6 hours at the beginning of the light phase. Fasting glucose was measured via tail cut using a hand-held glucometer (OneTouch Ultra2, LifeScan IP Holdings) with GenUltimate test strips (PharmaTech Solutions, Inc.). Mice received an intraperitoneal injection of either D-glucose (2g/kg lean mass) or insulin (1.0U/kg lean mass; Humulin R-500, Lilly) in PBS (for glucose and insulin tolerance tests, respectively), and the change in blood glucose concentration was monitored every 15 min for 2 hours.

### In vivo Induction of Lipolysis

At 22 weeks of age, mice were anesthetized with isofluorane and a blood sample was drawn via heparinized capillary from the retroorbital sinus. After a short recovery period, mice were injected intraperitoneally with 10 mg/kg isoproterenol (prepared in saline; #I6504, Sigma Aldrich). A second blood sample was collected from anesthetized mice 15 min post-injection. Blood was allowed to clot over ice before being centrifuged at 500g for 20 min at 4 °C. Glycerol and NEFA concentrations in serum were determined colorimetrically, using commercially available reagents (# F6428, Sigma Aldrich; NEFA-HR(2), Wako Life Sciences, Inc.).

### Tissue Collection

At 24 weeks of age mice were given a single intraperitoneal bolus of deuterium oxide (#151882, Sigma Aldrich) containing 0.9% NaCl, at a final enrichment of 4.5 ± 0.54% of total body water immediately prior to the light phase (ZT-0.5). All water sources, including sucrose drinking water, were removed from the cages at the time of injection. Seven hours later, mice were anesthetized with isofluorane and blood was drawn from the retroorbital sinus and allowed to clot over ice. An unclotted aliquot was snap frozen in liquid N_2_ for later determination of deuterium body water enrichment. Mice were then euthanized by cervical dislocation and tissues harvested. The left lateral lobe of the liver, right gonadal adipose depot (gWAT) and right dorsolumbar-inguinal adipose depot (iWAT) were immediately freeze-clamped between stainless steel paddles pre-cooled in liquid N_2_ and stored at −80°C for later analyses. The left iWAT depot was dissected out whole, quickly weighed, and placed into Krebs-Ringer-Bicarbonate-HEPES buffer (KRBH) for subsequent collagenase digestion and isolation of primary adipocytes. A portion of the median lobe of the liver was fixed in 10% neutral buffered formalin for 24 hours before being dehydrated in ethanol and imbedded for histology.

### Preparation of Adipocytes and ex vivo Lipolysis

Primary adipocytes were isolated from iWAT. Briefly, adipose tissue was allowed to recover in KRBH buffer containing 2.5% BSA for 20 min before being chopped into tiny pieces and incubated in KRBH buffer containing 0.5% BSA and 1% collagenase (type II; #17101015, Gibco) for one hour in a 37 °C shaking water bath. Primary adipocytes were separated from the stromal vascular fraction via a series of low centrifugation wash steps using KRBH containing 2.5% BSA. Aliquots of packed adipocytes were incubated in KRBH buffer with or without [100 μM] isoproterenol for 1 hour at 37 °C. Buffer infranatants were collected, heat inactivated for 10 min at 85 °C, and analyzed for glycerol content using a commercially available kit (#F6428, Sigma Aldrich). Adipocyte number was determined using a hemocytometer and cell counts were used to normalize rates of lipolysis to the number of cells present in the incubation medium.

### Tissue Metabolite Determination

Blood glucose concentrations were measured in whole-blood using a hand-held glucometer (OneTouch Ultra2, LifeScan IP Holdings) with GenUltimate test strips (PharmaTech Solutions, Inc.). Serum was separated from blood via centrifugation at 500g for 20 min at 4°C. Serum insulin and corticosterone concentrations were measured using ELISA’s (#90080 and #80556, Crystal Chemical). Liver triglycerides were extracted via chloroform-methanol (2:1) extraction, evaporated overnight, re-suspended in butanol-methanol-Triton-X114 and measured colorimetrically (#TR0100, Sigma Aldrich).

### De novo Fatty Acid and Triglyceride Synthesis

Deuterium oxide rapidly mixes with body water, allowing H^2^ to be incorporated into biosynthetic pathways that use H_2_O. Newly synthesized products can be quantified by measuring the H^2^ label present in the total pool of the product of interest. In this study, mice were provided a single bolus of 0.9% saline prepared with deuterium oxide seven hours before tissue collection and all other water sources were removed so as not to dilute the label. This short labelling period (hours rather than days) was chosen in order to minimize the contribution of recycled label (from product metabolism) into triglyceride-bound palmitate (which reflects the contribution of *de novo* fatty acid synthesis to the total triglyceride pool) and glycerol (which reflects the rate of triglyceride esterification) (Bederman et al., 2009; Brunengraber et al., 2003; Diraison et al., 1997).

Total triglycerides were isolated from liver samples via chemical hydrolysis and extraction (Brunengraber et al., 2003). Samples were dried down, derivatized, and converted to their trimethylsilyl derivatives for the measurement of glycerol and fatty acids (palmitate, oleate and stearate) after the addition of internal standards (Bederman et al., 2009). Isotope enrichments were determined by gas chromatography (GC)– mass spectrometry (MS) using chemical ionization. Deuterium labeling of body water was determined in whole-blood samples by acetone exchange (Bederman et al., 2009). All MS analyses were carried out on an Agilent 5977B MSD (CI or EI mode) coupled to a 7890B GC. The contribution of newly synthesized glycerol and fatty acids to the total triglyceride pool was calculated as previously described (Bederman et al., 2009; Brunengraber et al., 2003; Diraison et al., 1997).

### RNA isolation and preparation

RNA was extracted from liver and iWAT using TRI Reagent (#AM9738, Ambion) and column purified (#12183025 Purelink RNA mini kit, Life Technologies). RNA was either quantified spectrophotometrically (liver RNA; Nanodrop 2000, ThermoFisher Scientific) or by Agilent 2100 Bioanalyzer (iWAT RNA). cDNA was synthesized from purified liver RNA using a High-Capacity cDNA Reverse Transcription Kit (#4368813, Applied Biosystems). iWAT RNA was submitted to Novogene for cDNA library preparation and subsequent RNAseq analysis.

### q-PCR

Liver cDNA was combined with the appropriate working quantitative PCR master mix, containing Power SYBR Green (#4368708, Applied Biosystems) and the relevant primer pair (final concentration [100 nM] each; Integrated DNA Technologies). PCR conditions included an activation cycle of 95°C for 10 min followed by 45 amplification cycles of 15 s at 95°C, 15 s at 60°C, and 10 s at 73°C. CT values were quantified on QuantStudio 6 Flex Real-Time PCR System (Applied Biosystems). The ΔΔCT method was used to calculate the expression levels of mRNA transcripts using *Actb* as the housekeeping transcript. The expression of *Actb* in our liver samples was determined to be unaffected by sex or sucrose intake after being compared alongside other commonly used qPCR housekeeping transcripts, including *Rps18, Rpl13a*, *Rplp0, B2m* and *Gapdh*. Data were expressed relative to the female control group. The sequences of each of the primers used can be found in Supplementary File 1.

### cDNA Library Preparation and Transcriptomics

iWAT RNA samples were submitted to Novogene where mRNA was purified from total RNA using poly-T oligo-attached magnetic beads. After random fragmentation, NEB cDNA libraries were constructed, assessed for quality, and sequenced on an Illumina platform (paired-end 150bp). Downstream analysis was performed using a combination of programs including STAR, HTseq, Cufflink and Novogene’s wrapped scripts. Alignments were parsed using Tophat. Reference genome and gene model annotation files were downloaded from genome website browser (NCBI/UCSC/Ensembl) directly. Paired-end clean reads were aligned to the reference genome using STAR (v2.5). HTSeq v0.6.1 was used to count the read numbers mapped to each gene and the Fragments Per Kilobase of transcript per Million mapped reads (FPKM) of each gene was calculated based on the length of the gene and the mapped read counts.

### Differential Gene Expression and Gene Set Enrichment Analyses

FPKM data were compared in R Studio using the package DESeq2 v1.24.0(Love, 2014). Differential expression analyses were conducted to determine the effects of sucrose intake within each sex, the effect of sucrose intake with sex as a moderating effect, and to determine any sex:sucrose interactions. P-values were adjusted for multiple comparisons according to Benjamini and Hochberg(Benjamini Y, 1995), and transcripts with an adjusted p-value <0.05 were considered differentially expressed. String network analysis was performed on differentially expressed genes common to both male and female mice using string-db.org version 11.0(Szklarczyk et al., 2019). Gene Set Enrichment Analysis (GSEA) of all differentially expressed genes was completed in R Studio using the package fgsea v1.10.1(A, 2016). GSEA was completed with all gene sets in the Molecular Signatures Database reference gene set (MSigDB.v7.0), downloaded from https://www.gsea-msigdb.org/. A gene set was considered enriched when the nominal p-value was <0.01, the false discovery rate was <0.25, and the normalized enrichment score was >1.5 (for gene sets positively enriched) or less than −1.5 (for gene sets negatively enriched).

### Western blotting

Homogenates were prepared from ∼ 50 mg of frozen liver in RIPA buffer [Tris basic (50 mM), sodium deoxycholate (0.25%), NP-40 (1%), NaCl (150 mM), EDTA (1 mM), Na3VO4 (100 μM), NaF (5 mM), sodium pyrophosphate (10 mM), protease inhibitor cocktail] using stainless-steel beads and a Qiagen Tissue Lyser (30 Hz for 5 min). Homogenates were centrifuged at 4°C for 10 min at 14,000 g, after which the protein concentration of supernatants was determined by BCA assay. Lysates of equal protein concentration were prepared in 2× Laemmli buffer containing 2-mercaptoethanol and heated at 95°C for 5 min. Proteins were separated by SDS-PAGE and transferred to nitrocellulose membranes for Western blotting. Membranes were visualized for total protein using stain-free technology (BioRad) before being blocked in BSA for 1 h and incubated at 4°C overnight in the relevant primary antibody. Blots were visualized after a 1-h incubation with infrared anti-mouse or anti-rabbit secondary antibody, using a LI-COR Odyssey fluorescent Western blotting system. Protein expression was quantified using densitometry (Image Studio Lite; LI-COR) and normalized to total protein. b-Tubulin was included as a visual loading control for figures. Antibodies used include: SREBP-1c (#557036, BD Pharmingen), FAS (#3180 Cell Signaling Technology), ACC1/2 (#3676S, Cell Signaling Technology), phospho-ACC1/2 Ser79 (#3661, Cell Signaling Technology), b-Tubulin (#MA5-16308, Invitrogen), Goat anti-Rabbit IgG Alexa Fluor 680 secondary antibody (#A21109, Invitrogen) and Donkey anti-Mouse IgG Alexa Fluor 790 secondary antibody (#A11371, Invitrogen).

### Histology

Tissue samples were fixed in 10% neutral buffered formalin overnight, dehydrated in ethanol, cleared in Citrisolv and embedded in Paraplast plus. Sections were cut to 10μm and stained with hematoxylin and eosin. High-resolution images of slides were obtained at a 40X objective (EVOS XL, ThermoFisher Scientific).

### Microbiome Analysis With Illumina MiSeq Sequencing and Bioinformatics

Stool samples were collected fresh and snap frozen. Samples were resuspended in 500 μL of TNES buffer containing 20 μg of proteinase K and 150 μL of 0.1 zirconia beads. Following mechanical disruption using ultra-high-speed bead beating samples were incubated overnight at 55 °C with agitation. Total DNA was extracted using phenol chloroform isoamyl alcohol, and total DNA concentration per mg stool was determined by qRT-PCR. Purified DNA samples were sent to the Argonne National Laboratory (Lemont, IL) for amplicon sequencing using the NextGen Illumina MiSeq platform. Blank samples passed through the entire collection, extraction and amplification process remained free of DNA amplification.

Sequencing data were processed and analyzed using QIIME (Quantitative Insights into Microbial Ecology) 1.9.1. Sequences were first demultiplexed, then denoised and clustered into sequence variants. For bacteria we rarified to a depth of 10,000 sequences. Representative bacterial sequences were aligned via PyNAST, taxonomy assigned using the RDP Classifier against Greengenes (13.8) database. Processed data were then imported into Calypso 8.84 for further analysis and data visualization (Zakrzewski, 2017). For PICRUSt, biom tables were filtered against Greengenes databases, normalized, and sequences used to predict KEGG orthologs and pathway enrichments. The Shannon Index, Simpson Index, and Evenness were used to quantify alpha diversity (inter-sample) (Caporaso et al., 2010; Hughes, 2001). Bray-Curtis analysis was used to quantify beta diversity (intra-sample) (Lozupone et al., 2011), and the differences were compared using PERMANOVA, Anosim, and PERMDISP2. ANOVA, adjusted using the Bonferroni correction and false discovery rate for multiple comparisons (P<0.05), was used to quantify relative abundance of taxa among groups. Heatmaps were generated using Spearman’s rank correlation coefficient.

### Statistics

All statistical analyses were performed using R v3.6.1. running in R Studio v1.2.5033. Data with multiple repeated measures were analyzed by linear mixed effects modeling with likelihood ratio tests using the package lme4 (Bates D, 2015). All other data were assessed for homoscedasticity and normality using Levene’s tests (package: car (Fox J, 2019)) and Shapiro-Wilk tests, respectively. Any heteroscedastic data sets were log-transformed to attain homoscedasticity and re-assessed for normality. Normal data were analyzed using two-way ANOVA with interaction models and Tukey post-hoc analysis. Any two-way factorial non-normal data were analyzed using Scheirer-Ray-Hare tests (package: rcompanion (Mangiafico, 2016)), with post hoc Wilcoxon Rank Sum test adjusted for multiple comparisons using the Benjamini and Hochburg correction. Only the results of significant sex:sucrose interactions are reported. Significance was set *a priori* with a p-value <0.05 considered significant.

## Results

### Sucrose Intake Increases Adiposity, with Males Having a More Robust Response than Females

We *ad libitum* fed male and female mice standard rodent chow (3.1 kCal/g, 17% kCal from fat, 58% from carbohydrate, 25% from protein) with water from weaning until 10 weeks of age, after which mice continued to receive the chow diet with water (control groups) or chow with water containing 10% w/v sucrose (treatment groups) for 12 weeks.

Prior to the treatment period (4-10 weeks of age, Figure 1A, unshaded area), we observed a main effect of sex on body weight (^†^p=3.14^−11^) and lean mass gains (^†^p=8.62^−14^), but not fat mass gains (p=0.507). Treatment designation had no effect on any of the body composition parameters (p=0.283 for body weight, p=0.642 for body fat and p=0.383 for lean mass).

**Figure 1:**
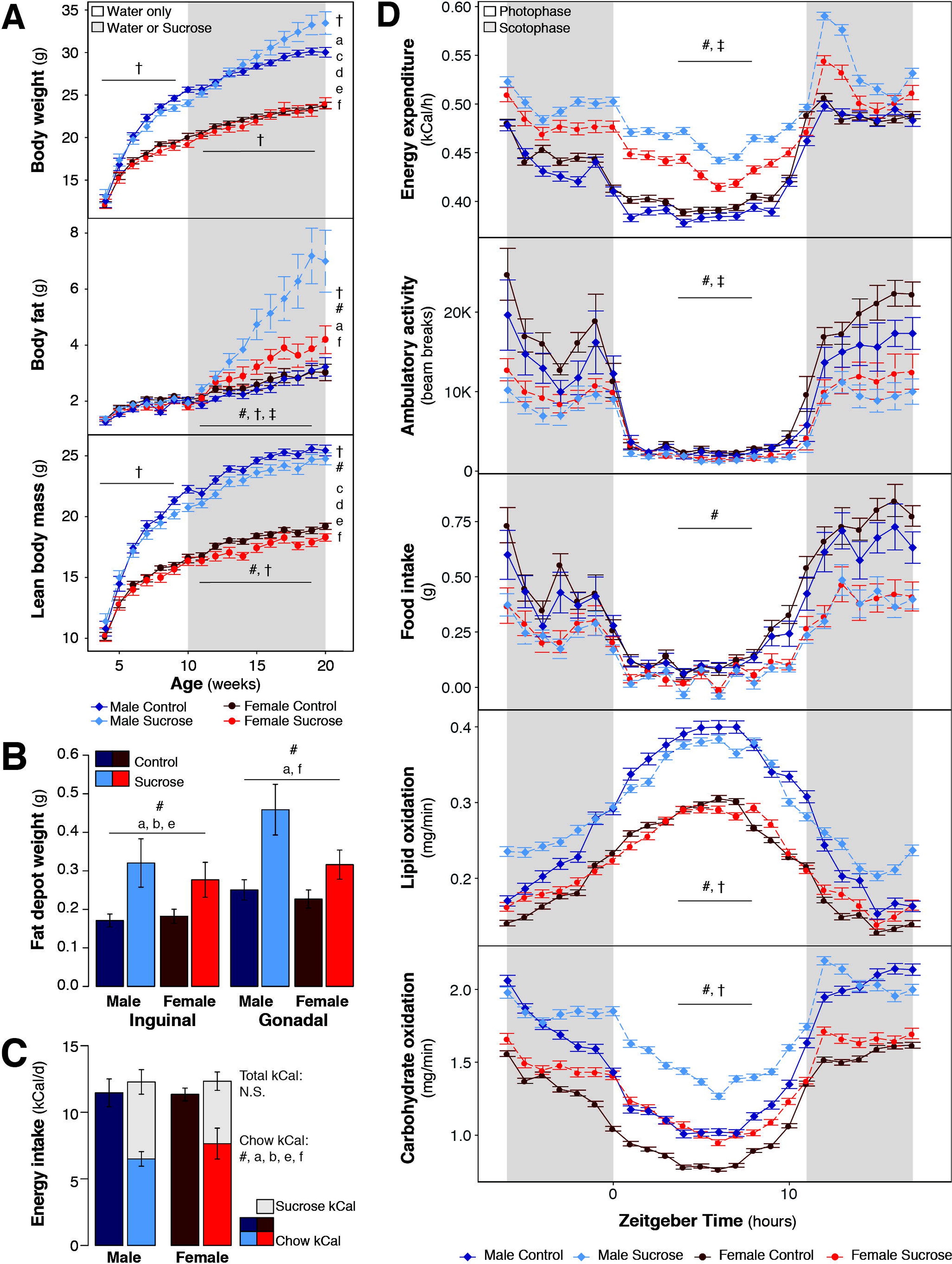
Chronic liquid sucrose intake leads to reorganization of the body’s energy stores in favor of increased adiposity and this effect is greater in male mice. During the treatment period there was an effect of sex on total body weight (upper panel, A), body fat (middle panel, A) and lean mass (lower panel, A; all n=13-18/group). Sucrose intake increased both body fat and lean mass, and a sex:sucrose interaction effect was observed for body fat. In line with an increase in total body fat, sucrose intake also led to increased weight of individual adipose depots (B; n=13-17/group) despite energy intake being similar across groups (C; n=7-13/group). Food intake was reduced in the groups receiving sucrose (C and middle panel, D; n=7-14/group), and energy expenditure was increased (upper panel, D; n=7-14/group). Sucrose intake also decreased ambulatory activity (second panel, D; n=7-14/group) and increased both lipid and carbohydrate oxidation (lower two panels, D; n=7-14/group). Both lipid and carbohydrate oxidation were also modified by sex, whereas sex:sucrose interaction effects were observed for both energy expenditure and ambulatory activity. Data presented are means ± se. In panels C and D, data are the group means determined from the mean values for each individual mouse continuously measured daily for a minimum of 7 d. Symbols denote p<0.05, where # indicates a sucrose intake effect, † indicates an effect of sex, and ‡ indicates a sex:sucrose interaction. For post-hoc analyses, a=Male Control vs Male Sucrose, b=Female Control vs Female Sucrose, c=Male Control vs Female Control, d=Male Sucrose vs Female Sucrose, e=Male Control vs Female Sucrose, f=Male Sucrose vs Female Control. In panel A, post-hoc analysis was only completed on data collected at 20-weeks of age.

During the treatment period (10 weeks of age onward; Figure 1A, shaded area), sucrose intake resulted in increased body fat (^#^p=3.7^−4^) and decreased lean mass (^#^p=0.012), whereas no effect of sucrose was observed for total body weight (p=0.325). We observed a sex:sucrose interaction for body fat (^‡^p=0.033). By 20 weeks of age, sucrose intake had increased body weight in male (98.1%, ^a^p=0.049) but not female mice (p=0.75). Increased weight in males following sucrose intake was associated with an increase in body fat (445.6%, ^a^p=9.7^−4^). We continued to observe an effect of sex on body weight (^†^p<1.0^−15^) and lean mass (^†^p<1.0^−15^) during this time, whereas there was also an effect of sex on body fat (^†^p=0.048).

To determine the effect of sucrose intake on specific adipose tissue depots, we carefully dissected out and weighed both subcutaneous (iWAT) and visceral (gWAT) depots (Figure 1B). No effect of sex was observed for either inguinal (p=0.731) or gonadal (p=0.084) adipose tissue, although we did detect an effect of sucrose intake (inguinal: ^#^p=6.6^−4^, gonadal: ^#^p=5.5^−4^). In both males and females, sucrose intake increased the mass of inguinal fat (88.2%, ^a^p=0.045, and 55.6%, ^b^p=0.049, respectively). Sucrose intake also increased gonadal fat mass in males (84.0%, ^a^p=0.008), although in females the increase in gonadal fat mass did not attain statistical significance (49.1%, p=0.085). Together, these results demonstrate that chronic sucrose intake increases adiposity and that the effect of sucrose on adiposity gains is similar in males and females.

### Chronic Liquid Sucrose Intake Modifies Energy Balance

To determine the role of energy intake in determining adiposity gains following chronic sucrose intake, we measured the average daily food and water intake of mice individually housed between weeks 10-11 of treatment (Figure 1C and middle panel of 1D). In both sexes, sucrose intake led to reduced energy intake from food (Figure 1C; ^#^p=1.8^−4^; −43.3% in males, ^a^p=0.014; −32.6% in females, ^b^p=0.046) and, as a result, total energy intake was similar between all four groups, regardless of sex or treatment designation (Figure 1C). Notably, absolute fluid intake was markedly elevated in response to sucrose intake (^#^p=8.5^−5^) but not sex (p=0.273), with increased intake of fluid in both male (194.9%, ^a^p=0.005) and female (182.9%, ^b^p=0.039) mice if sucrose was present in the drinking water.

To determine the relative contribution of energy expenditure to adiposity gains following liquid sucrose intake, we performed indirect calorimetry and activity monitoring experiments in parallel with measurements of energy intake beginning at 20 weeks of age (10 weeks of sucrose treatment). Sex had no effect on energy expenditure (p=0.627), whereas sucrose intake increased energy expenditure in both sexes (Figure 1D, upper panel; ^#^p=3.9^−5^). A sex:sucrose interaction was also observed (^‡^p=0.025), with males appearing to have a greater increase in energy expenditure than females after sucrose intake. Changes in physical activity did not explain the increase in energy expenditure observed following sucrose intake, as ambulatory activity was decreased in both sexes in response to sucrose intake (^#^p=1.2^−4^), primarily during the scotophase (shaded area of Figure 1D), with a sex:sucrose interaction also observed (^‡^p=0.038). Similar results were found for stereotypic activity (not shown). To establish whether changes in energy substrate oxidation might be contributing to sucrose-induced increases in adiposity, we calculated the rates of both lipid (fourth panel of Figure 1D) and carbohydrate oxidation (last panel of Figure 1D), observing effects of both sex (^†^p=8.1^−5^ and ^†^p=9.2^−4^, respectively) and sucrose intake (^#^p=2.8^−4^ and ^#^p=0.002, respectively), with males appearing to have greater magnitude increases than females following sucrose intake. To establish the effects of sucrose intake on energy balance in the absence of thermal stress, we repeated these experiments with animals housed at the lower end of their thermoneutral zone (28°C; Supplementary Figure 1). An effect of sex was observed for energy expenditure (^†^p=0.006), food intake (^†^p=1.1^−4^), ambulatory activity (^†^p=1.0^−5^) and both lipid (^†^p=0.027) and carbohydrate oxidation (^†^p=2.4^−8^). Similar to our findings at 25°C, sucrose intake under thermoneutral conditions increased energy expenditure (^#^p=7.7^−4^) and lipid oxidation (^#^p=0.007), while decreasing food intake (^#^p=0.006). In the absence of thermal stress, sucrose intake reduced carbohydrate oxidation (^#^p=2.4^−8^), whereas ambulatory activity was unaffected (p=0.350).

To determine whether impairments in glucose uptake or insulin responsiveness were contributing to sucrose-induced changes in substrate utilization we performed insulin and glucose tolerance tests (Supplemental Figure 2A-D). While an effect of sex was observed for both tests (^†^p=0.003 for insulin tolerance test, ^†^p=7.4^−4^ for glucose tolerance test), sucrose intake had no effect. Neither sex nor sucrose intake affected fasting blood glucose concentrations (Supplemental Figure 2E), whereas there was an effect of sex on fasting serum insulin (Supplemental Figure 2F; ^†^p=1.1^−4^), with 26.5% lower concentrations observed in the female control group compared to the male control group (^c^p=7.4^−4^). Serum corticosterone was similar across all four groups (Supplemental Figure 2G).

Taken together, our findings for energy balance and glucose homeostasis suggest there may be a disconnect between energy intake, energy expenditure and adiposity that occurs in the absence of systemic insulin resistance following chronic liquid sucrose intake.

### Chronic Liquid Sucrose Intake Increases Hepatic Triglyceride Content via Sex-Specific Mechanisms

We first confirmed that chronic liquid sucrose intake at physiologically relevant concentrations resulted in hepatic steatosis by quantifying liver triglyceride content biochemically (Figure 2A) and visualizing liver histology (Figure 2B; representative image after staining with hematoxylin and eosin). Effects of both sex (^†^p=1.3^−7^) and sucrose intake (^#^p=4.2^−9^) were observed for total triglyceride content. Sucrose intake increased liver triglycerides in both males (271.0%, ^a^p=4.3^−5^) and females (434.6%, ^b^p=2.6^−6^) compared to their corresponding control groups, with female mice in the sucrose group having an exacerbated response compared to males (266.1%, ^d^p=2.3^−5^).

**Figure 2:**
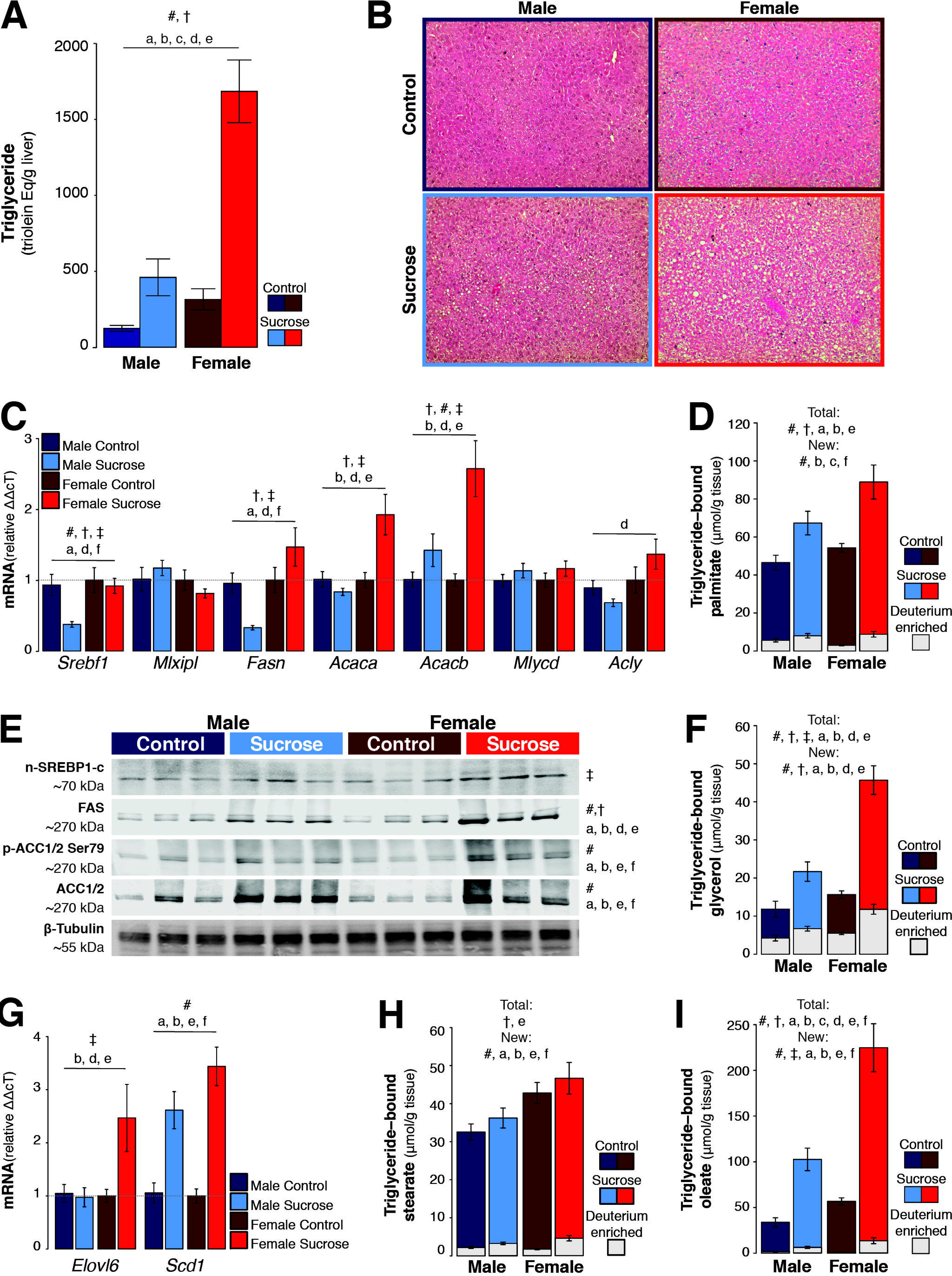
Chronic liquid sucrose intake increases hepatic triglyceride content via sex-specific mechanisms. Hepatic triglyceride content was increased in both sexes following chronic liquid sucrose intake; however, the magnitude of the sucrose effect was greater in female mice (A, quantification from biochemical analysis, n=11-14/group; and B, representative H&E stain). Transcriptional regulation of fatty acid synthesis (C; n=11-12/group) also exhibited sex-specific effects of sucrose intake, with male mice downregulating transcripts for *Srebf1* and *Fasn*, and females upregulating transcripts for *Acaca* and *Acacb*. Transcriptional data were supported by mass spectrometry data which demonstrate that although sucrose intake increased the total pool of triglyceride-bound palmitate in both sexes, the pool of deuterium-enriched palmitate bound to triglyceride (the fraction derived from *de novo* fatty acid synthesis) was only increased in female mice after sucrose intake (D; n=11-14/group). The abundance of proteins encoded by the transcripts differentially expressed between males and females in response to sucrose intake also demonstrated sex-dependent effects; particularly FAS which was increased to a greater magnitude in females after sucrose intake, while both phosphorylated (inhibitory) and total ACC/1/2 showed sucrose-induced increases only. A sex:sucrose interaction was observed for nSREBP1-c (E; quantification from n=6/group, n=3/group shown). We measured the amount of total and newly synthesized triglyceride-bound glycerol, which represent the total pool of hepatic triglyceride and the triglyceride pool that is newly esterified, respectively. Both total glycerol and glycerol from glyceroneogenesis were increased in both sexes in response to sucrose intake, although the magnitude of these increases was greater in female mice (F; n=11-14/group), a finding consistent with the results in panel D. Transcriptional regulation of fatty acid modifications also exhibited sex-specific sucrose effects (G; n=11-12/group). Only female mice upregulated the fatty acid elongation transcript *Elovl6,* whereas both sexes upregulated the desaturase transcript *Scd1* following sucrose intake. This, too, was supported by mass spectrometry data, demonstrating a sex effect for total triglyceride-bound stearate, whereas sucrose intake increased newly elongated stearate in both sexes (H; 11-14/group). Both effects of sex and sucrose intake were observed for total triglyceride-bound oleate, with females having more triglyceride-bound oleate than males under non-sucrose conditions as well as a greater magnitude increase in triglyceride-bound oleate following sucrose intake. Newly desaturated triglyceride-bound oleate was similarly increased in response to sucrose intake, with a sex:sucrose interaction effect also observed (I; n=11-14/group). Data presented are means ± se. Symbols denote p<0.05, where # indicates a sucrose intake effect, † indicates an effect of sex, and ‡ indicates a sex:sucrose interaction. For post-hoc analyses, a=Male Control vs Male Sucrose, b=Female Control vs Female Sucrose, c=Male Control vs Female Control, d=Male Sucrose vs Female Sucrose, e=Male Control vs Female Sucrose, f=Male Sucrose vs Female Control.

To identify what might be causing these sex differences at the transcriptional level, we measured the expression of a selection of gene transcripts known to regulate *de novo* fatty acid synthesis (Figure 2C). An effect of sex was observed for several transcripts, including: *Srebf1* (^†^p=0.01), *Fasn* (^†^p=5.7^−4^), *Acaca* (^†^p=0.002) and *Acacb* (^†^p=0.050). An effect of sucrose intake was observed for *Srebf1* (^#^p=0.010) and *Acacb* (^#^p=3.6^−4^), while the effect of sucrose intake did not attain statistical significance for *Fasn* (p=0.055) or *Acaca* (p=0.056). In response to sucrose intake, males had reduced expression of *Srebf1* and *Fasn* (59.4%, ^a^p=0.003 and 65.4%, ^a^p=0.001; respectively), whereas females had increased expression of *Acaca* and *Acacb* (92.2%, ^b^p=0.002 and 157.1%, ^b^p=7.1^−4^; respectively). Sex:sucrose interactions were observed for *Srebf1* (^‡^p=0.008), *Fasn* (^‡^p=4.4^−4^), *Acaca* (^‡^p=0.001) and *Acacb* (^‡^p=0.040). *Acly* also showed a tendency toward a sex:sucrose interaction, although this did not attain statistical significance (p=0.059).

Since the transcript data suggested that differences in *de novo* fatty acid synthesis may be driving some of the sex differences we observed in response to sucrose intake, we used a deuterium enrichment strategy to measure the amount of newly synthesized palmitate in the hepatic triglyceride pool (Figure 2D). We observed both sex (^†^p=0.013) and sucrose intake effects (^#^p=5.2^−5^) for total triglyceride-bound palmitate (Figure 2D, colored bars). Both males and females had increased total triglyceride-bound palmitate following sucrose intake compared to their corresponding control groups (44.7%, ^a^p=0.03 and 64.9%, ^b^p=0.01; respectively). Of the total pool of triglyceride-bound palmitate, the contribution made by newly synthesized palmitate (Figure 2D, grey inlaid bars) showed an effect of sucrose intake (^#^p=1.1^−4^), although newly synthesized palmitate was only increased in female mice (66.4%, ^b^p=9.2^−5^).

Given that *de novo* fatty acid synthesis can be regulated post-transcriptionally, we also measured the relative abundance of proteins encoded by the transcripts we identified as being differentially expressed between male and female mice following sucrose intake (Figure 2E). FAS was the only protein we measured that demonstrated an effect of sex (^†^p=0.001), although the ratio of phosphorylated ACC1/2 (inhibitory phosphorylation) at Ser79 to total ACC also demonstrated an effect of sex (^†^p=0.034), while there was a sex:sucrose interaction for the abundance of nuclear SREBP1-c (^‡^p=0.046). An effect of sucrose intake was observed for FAS (^#^p=1.0^−6^), phospho-ACC1/2 at Ser79 (^#^p=1.2^−5^) and total ACC1/2 (^#^p=7.3^−6^), with each of these proteins being increased in abundance in response to sucrose intake in both male (98.0%, ^a^p=0.01; 223.0%, ^a^p=0.001 and 178.4%, ^a^p=0.002; respectively) and female mice (129.8%, ^b^p=2.2^−5^; 225.3%, ^b^p=0.007 and 223.1%, ^b^p=0.003; respectively). The sucrose-induced increase in hepatic FAS protein was also greater in female compared to male mice (57.2%, ^d^p=0.003). Taken together, these data provide mechanistic insight into how chronic sucrose intake results in hepatic steatosis and identifies sex as a moderating factor that can determine the contribution of *de novo* fatty acid synthesis to the hepatic triglyceride pool. Specifically, we show that female mice have increased rates of *de novo* fatty acid synthesis which is regulated, at least in part, by *Srebf1*/SERBP1-c and the transcription factor’s targets, *Fasn*/FAS and *Acaca*/*Acacb*.

Since upregulation of *de novo* fatty acid synthesis was not the main factor driving the increase in liver triglyceride we observed following chronic liquid sucrose intake, we also determined the relative contribution made by fatty acid re-esterification by measuring glyceroneogenesis via deuterium incorporation into triglyceride-bound glycerol (Figure 2F). We observed effects of sex (^†^p=2.1^−6^), sucrose intake (^#^p=9.2^−10^) and a sex:sucrose interaction (^‡^p=0.007) for the total pool of triglyceride-bound glycerol (Figure 2F, colored bars). Both male (83.0%, ^a^p=0.005) and female (192.1%, ^b^p=6.1^−8^) mice receiving sucrose had an increase in the total pool of triglyceride-bound glycerol compared to their respective control groups, whereas female mice receiving sucrose had more glycerol than male mice receiving sucrose (110.7%, ^d^p=2.5^−6^). Of the total pool of triglyceride-bound glycerol, the contribution made by newly synthesized glycerol (Figure 2F, grey inlaid bars) was dependent on both sex (^†^p=3.7^−4^) and sucrose intake (^#^p=4.0^−6^). Similar to our observations for the total glycerol pool, newly synthesized glycerol increased in response to sucrose intake in both males (58.7%, ^a^p=0.035) and females (114.0%, ^b^p=4.9^−4^), and this effect was greater in female mice receiving sucrose compared to males (75.6%, ^d^p=0.003). These findings suggest that hepatic triglyceride accumulation following chronic liquid sucrose intake is primarily driven by the re-esterification of fatty acids derived from extrahepatic sources in male mice, and by both *de novo* fatty acid synthesis and re-esterification of extrahepatic fatty acids in female mice.

After identifying an effect of sucrose intake for the hepatic expression of the transcripts *Elovl6* and *Scd1* (Figure 2G), we leveraged the presence of deuterium labelling in the liver samples to look at the effect of sex and sucrose intake on fatty acid modifications and their contribution to the hepatic triglyceride pool, including elongation of palmitate into stearate, and desaturation of stearate into oleate (Figure 2H-I). Total triglyceride-bound stearate was affected by sex (Figure 2H, colored bars; ^†^p=8.8^−4^), whereas newly elongated stearate was increased by sucrose intake only (Figure 2H, grey inlaid bars; ^#^p=2.5^−6^), with increases observed in both males (37.4%, ^a^p=0.02) and females (164.4%, ^b^p=1.0^−5^) in response to sucrose intake. Total triglyceride-bound oleate was affected by both sex (Figure 2I, colored bars; ^†^p=5.4^−7^) and sucrose intake (^#^p=1.8^−13^), with female mice having more total oleate than male mice in both the control (68.0%, ^c^p=0.007) and sucrose groups (119.1%, ^d^p=1.1^−4^), and sucrose intake increasing total oleate in both male (204.1%, ^a^p=1.0^−7^) and female mice (296.5%, ^b^p=3.5^−8^). We observed an effect of sucrose intake on newly desaturated triglyceride-bound oleate (Figure 2I, grey inlaid bars; ^#^p=1.0^−12^), as well as a sex:sucrose interaction (^‡^p =0.008). Relative to the control groups, sucrose intake increased newly desaturated oleate in both male (297.2%, ^a^p=2.1^−5^) and female mice (1,382.9%, ^b^p=3.2^−10^). Together, these data suggest both sex and sucrose intake moderate the modification of fatty acids and their incorporation into the hepatic triglyceride pool.

### Sucrose Intake Potentiates β-Adrenergic Receptor-Stimulated Increases in Serum Glycerol Concentrations

To determine if increased availability of fatty acids from extrahepatic sources was contributing to the increase in re-esterification and the subsequent steatosis we observed in response to chronic liquid sucrose intake, we measured serum glycerol and NEFA in mice both before (basal) and 15 min after stimulating lipolysis *in vivo* with the non-specific b-adrenergic receptor (β-AR) agonist isoproterenol (stimulated; Figure 3A). For serum glycerol (representative of systemic lipolysis; upper panels, Figure 3A), we observed effects of stimulation (^§^p<1.0^−15^), sex (^†^p=3.7^−8^), sucrose intake (^#^p=1.7^−8^) and a stimulation:sucrose interaction (∞p=0.031). Under basal conditions, there was an effect of sucrose intake in female mice, with females having 73.2% greater glycerol concentrations than males after sucrose intake (^d^p=0.039) and 66.2% greater glycerol concentrations compared to the female control group (although, the latter did not attain statistical significance, p=0.054). After β-AR stimulation, sucrose intake potentiated the increase in glycerol in both males (34.8%, ^a^p=0.007) and females (53.5%, ^b^p=1.6^−4^), with greater glycerol concentrations being observed in females compared to males following sucrose intake (37.0%, ^d^p=3.2^−4^). For serum NEFA (representative of both systemic lipolysis and fatty acid clearance/tissue uptake; lower panels, Figure 3A), we observed effects of stimulation (^§^p<1.0^−15^), sex (^†^p=0.021) and a sex:sucrose interaction (^‡^p=0.006). No differences were observed in basal NEFA levels; however, after β-AR stimulation, sucrose intake potentiated the increase in serum NEFA in females when compared to males (18.3%, ^d^p=0.030).

**Figure 3:**
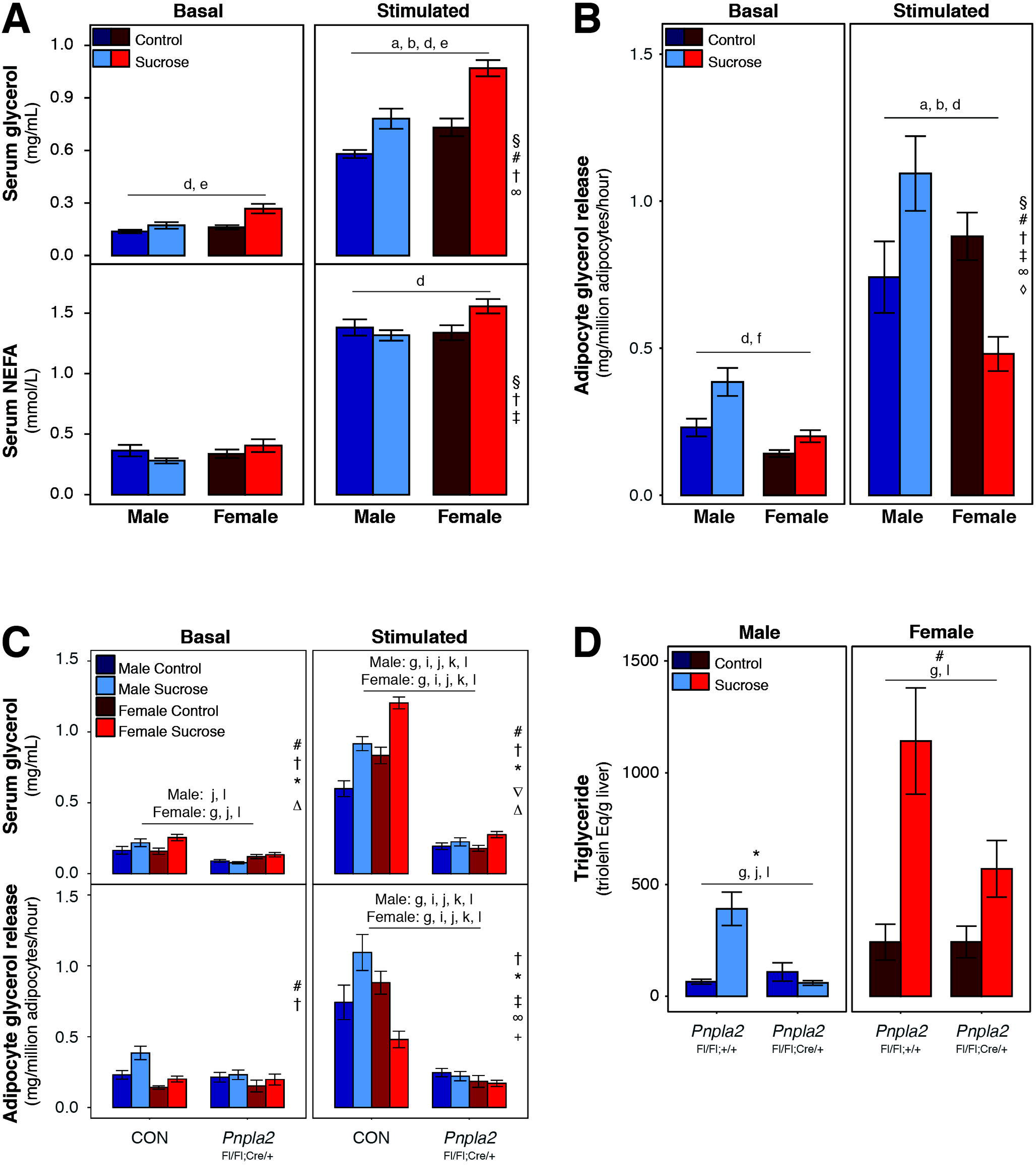
Impairing adipocyte lipolysis protects against sucrose-induced hepatic steatosis. Serum glycerol appearance in response to *in vivo* β-AR stimulation with isoproterenol was increased in both sexes following chronic liquid sucrose intake, with potentiation of stimulated glycerol appearance being greatest in female mice (A, upper panels; n=14-18/group). A sex:sucrose interaction effect was observed for the serum appearance of non-esterified fatty acids (NEFA; A, lower panels; n=14-18/group). In isolated adipocytes, sucrose intake resulted in potentiation of stimulated glycerol release only in adipocytes from male mice, with sucrose intake attenuating the induction of stimulated glycerol release in adipocytes from female mice compared to their respective control group (B; n=16-21/group). Compared to mice with control genotypes (CON), β-AR stimulation did not potentiate release of glycerol in either serum (C, upper panels) or isolated adipocytes (C, lower panels) from mice with adiponectin-driven knock-out of adipose triglyceride lipase (*Pnpla2*^Fl/Fl^;Cre^/+^; C; n=5-7/group). The blunted adipocyte lipolysis response in knockout mice prevented (males) or attenuated (females) sucrose-associated hepatic triglyceride accumulation (D; n=5-12/group). Data presented are means ± se. Symbols denote p<0.05, where § indicates an effect of stimulation, # indicates a sucrose intake effect, † indicates an effect of sex, ‡ indicates a sex:sucrose interaction, ∞ indicates a sucrose: stimulation interaction, and ◊ indicates a sex:sucrose:stimulation interaction and * indicates a genotype:sucrose interaction. For post-hoc analyses, a=Male Control vs Male Sucrose, b=Female Control vs Female Sucrose, c=Male Control vs Female Control, d=Male Sucrose vs Female Sucrose, e=Male Control vs Female Sucrose, f=Male Sucrose vs Female Control, g=CON or *Pnpla2*^Fl/Fl;+/+^ Control vs CON or *Pnpla2*^Fl/Fl;+/+^ Sucrose, h= *Pnpla2*^Fl/Fl;^Cre/^+^ Control vs *Pnpla2*^Fl/Fl;^Cre/^+^ Sucrose, i=CON or *Pnpla2*^Fl/Fl;+/+^ Control vs *Pnpla2*^Fl/Fl;^Cre/^+^ Control, j=CON or *Pnpla2*^Fl/Fl;+/+^ Sucrose vs *Pnpla2*^Fl/Fl;^Cre/^+^ Sucrose, k=CON or *Pnpla2*^Fl/Fl;+/+^ Control vs *Pnpla2*^Fl/Fl;^Cre/^+^ Sucrose, l=CON or *Pnpla2*^Fl/Fl;+/+^ Sucrose vs *Pnpla2*^Fl/Fl^;Cre/^+^ Control.

### Sex Moderates the Adipocyte Lipolysis Response to β-AR Stimulation After Chronic Sucrose Intake

To test whether adipocyte lipolysis was responsible for the potentiated induction of serum glycerol appearance we observed following systemic β-AR stimulation, we isolated mature adipocytes from iWAT and measured the rate of glycerol release under basal or isoproterenol-stimulated conditions (Figure 3B). We observed effects of stimulation (^§^p<1.0-5), sex (^†^p=7.0^−5^) and sucrose intake (^#^p=0.008). We also observed sex:sucrose (^‡^p=2.9^−4^), stimulation:sucrose (∞p=0.036) and stimulation:sex:sucrose interaction effects (^◊^p=0.005). Under basal conditions, males had increased lipolysis compared to females after sucrose intake (61.5%, ^d^p=0.009). It also appeared that males receiving sucrose had increased basal lipolysis compared to the male control group; however, this did not attain statistical significance (61.6%, p=0.052). Under stimulated conditions, sucrose intake potentiated the effect of β-AR stimulation on lipolysis in adipocytes from male mice (47.5%, ^a^p=0.026), whereas it attenuated the increase in lipolysis in adipocytes from female mice (−52.0%, ^b^p=0.031). Stimulated lipolysis was also 127.8% greater in males than females after sucrose intake (^d^p=6.0^−4^). These findings suggest that in male mice, sucrose intake leads to increased adipocyte lipolysis via adipocyte-autonomous mechanisms whereas the potentiation of systemic lipolysis in females likely requires a non-adipocyte intermediate to facilitate the potentiation effect of sucrose intake on β-AR-dependent lipolysis *in vivo*.

### Selective Inhibition of Adipocyte Lipolysis Protects Against Sucrose-Induced Hepatic Steatosis

To confirm whether increased adipocyte lipolysis contributes to the development of hepatic steatosis following chronic liquid sucrose intake, we repeated our sucrose drinking paradigm in mice homozygous for a LoxP-modified *Pnpla2* allele (Sitnick et al., 2013) with or without hemizygosity for an adipose tissuespecific Cre recombinase (*Adipoq*) (Eguchi et al., 2011). *Pnpla2* encodes adipose triglyceride lipase (ATGL), the rate-limiting enzyme of lipolysis. Mice missing ATGL from their adipose tissue had a marked impairment in β-AR-stimulated serum glycerol appearance (upper panel, Figure 3C), and the β-AR stimulated induction of lipolysis in isolated adipocytes from iWAT was almost completely abolished (lower panel, Figure 3C), suggesting that unimpaired adipocyte lipolysis is required for *in vivo* induction of lipolysis via β-AR stimulation, as well as sucrose-induced potentiation of the lipolysis response to β-AR stimulation. Our data also show that unimpaired adipocyte lipolysis is necessary for sucrose-mediated hepatic steatosis (Figure 3D). In male adipose-specific ATGL knockout mice (*Pnpla2*^Fl/Fl^Cre/^+^) and their control genotype counterparts (*Pnpla2*^Fl/Fl;+/+^) we did not observe significant effects of genotype (p=0.108) or sucrose intake (p=0.066) but we did observe a genotype:sucrose interaction effect (*p=0.017). This effect was associated with an increase in liver triglyceride content in mice with the control genotype after sucrose intake relative to the non-sucrose group (499.7%, ^g^p=0.004), whereas we observed no difference in knockout males after sucrose intake (p=0.999). Liver triglyceride content was also increased in male mice with the control genotype after sucrose intake compared to knockout males that also received sucrose, (556.7%, ^j^p=0.005). In female mice, we observed an effect of sucrose intake (^#^p=6.9^−5^) but not genotype (p=0.463). Similar to our observations in male mice, sucrose intake increased liver triglyceride content in female mice with the control genotype relative to the non-sucrose mice with the control genotype (371.5%, ^g^p=0.002). In female mice with adipose-specific ATGL knockout, we observed no significant difference between the sucrose and non-sucrose groups (p=0.151), nor did we see a difference between knockout females and females with the control genotype after sucrose intake (p=0.158), suggesting an intermediate effect of ATGL knockout on liver triglyceride content in female mice. Taken together, these data demonstrate that in male mice, adipocyte lipolysis is necessary for the development of hepatic steatosis following chronic liquid sucrose intake, whereas in female mice, impairment of adipocyte lipolysis attenuates but does not completely prevent hepatic triglyceride accumulation. These findings are in line with our earlier observation that in addition to increased NEFA re-esterification following sucrose intake (Figure 2F), female mice have increased rates of *de novo* fatty acid synthesis (Figure 2D), which would be expected to contribute to liver triglyceride content independently of NEFA derived from adipocyte lipolysis.

### Sucrose Intake Elicits Distinct Transcriptional Responses in Male and Female Adipose Tissue

To gain insight into the adipose-specific mechanisms contributing to the divergent lipolysis responses we observed in adipocytes from male and female mice after chronic sucrose intake (Figure 3B), we sought to identify how liquid sucrose consumption altered the adipose tissue transcriptome (Figure 4). Compared to their respective control groups, sucrose intake led to upregulation of 1045 gene transcripts in female mice and 128 gene transcripts in male mice (Figure 4A and Supplementary File 2). Only 6 transcripts were commonly upregulated by chronic sucrose intake in both sexes; these were: *Tma7*, *Ctsc*, *Rab21, Skap2, Tmf1* and *Pla2g7* (Figure 4B). We observed 1261 gene transcripts downregulated in adipose tissue from female mice after sucrose intake, whereas 151 transcripts were downregulated in male adipose tissue (Figure 4A and Supplementary File 2). Ninety-one transcripts were commonly downregulated in both male and female mice following sucrose intake (Figure 4B).

**Figure 4:**
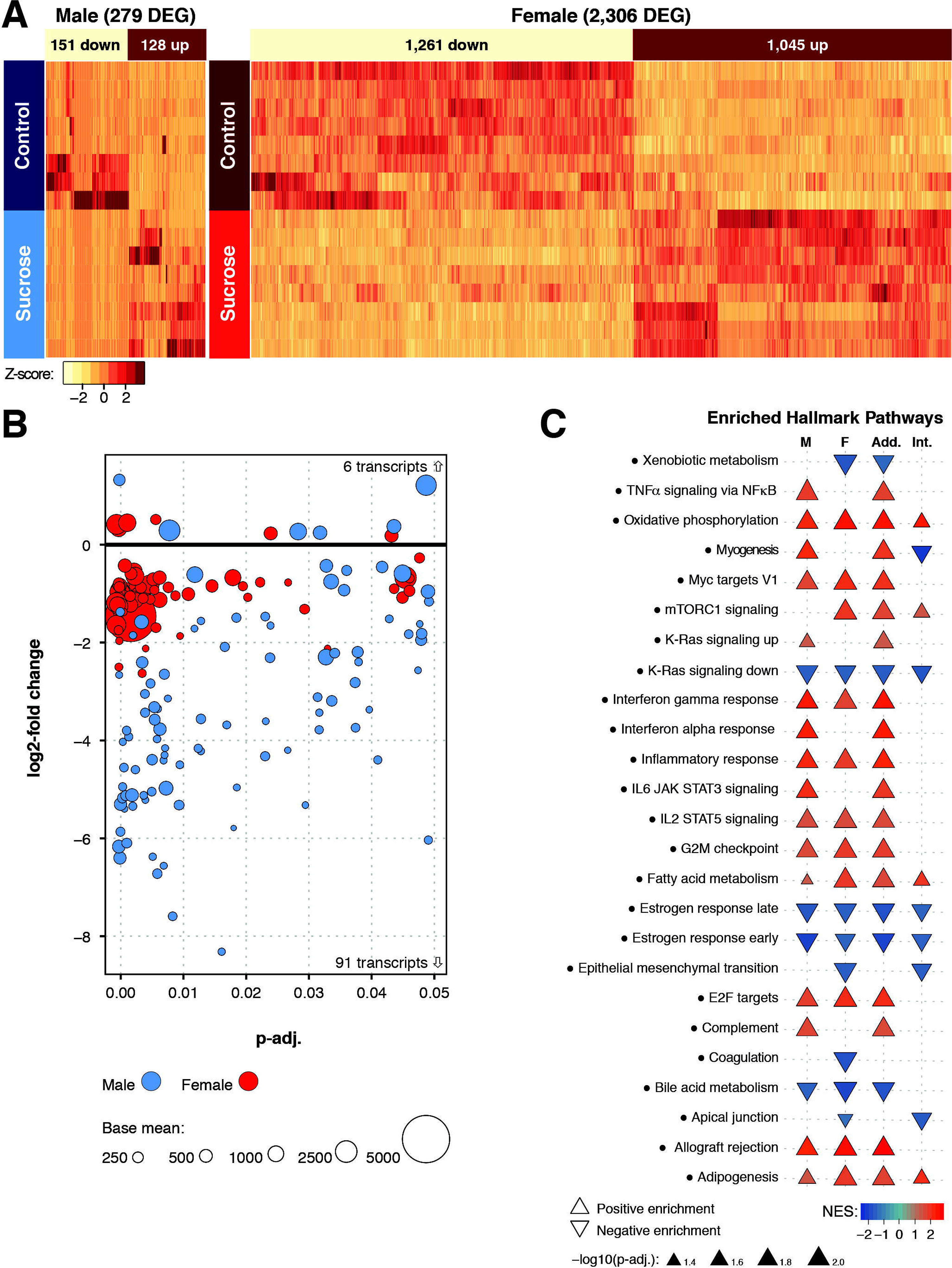
Sucrose intake elicits distinct transcriptional responses in adipose tissue. Heat maps demonstrate a marked difference in the number of transcripts differentially expressed in inguinal adipose tissue following chronic liquid sucrose intake in each sex, as determined by RNAseq (A; n=8/group). There were only 6 upregulated and 91 downregulated DEG common to both sexes (B). Top 25 gene sets enriched according to the Hallmark database (C; see also: suppl. file. 4). NES, normalized enrichment score; M, male; F, female; Add. additive effect of sex on the effect of sucrose intake; Int. sex:sucrose interaction effect.

To tease out any functional relationships between the differentially expressed genes common to both male and female mice, we performed string network analysis (Supplementary Figure 3A). The only functional relationship identified for a protein encoded by one of the commonly upregulated transcripts was *Rab21*, which formed an interaction network with other Rab proteins encoded by downregulated transcripts: *Rab17* and *Rab25*. Several larger interaction networks were observed for proteins encoded by the transcripts commonly downregulated. These networks contributed to functional enrichment of pathways involved in tissue morphogenesis, cell proliferation and differentiation, as well as cell-cell adhesion and cell-cell communication (Supplementary File 3).

To identify which pathways and networks altered by sucrose intake, including which pathways were also influenced by sex, we performed gene-set enrichment analyses comparing all of the differentially expressed genes we identified with the reference gene sets found in the Molecular Signatures Database (Supplementary File 4). We found 2048 pathways enriched in female mice following sucrose intake (1496 positively enriched, 552 negatively enriched), and 1378 pathways enriched in male mice (1168 positively enriched, 210 negatively enriched). An additive effect of sex on sucrose intake was observed for 2049 pathways (1650 positively enriched, 399 negatively enriched), whereas a sex:sucrose interaction effect was observed for 591 pathways (131 positively enriched, 460 negatively enriched). Among the positively enriched gene sets, those representing mitochondrial biogenesis, aerobic metabolism, and inflammation and immune responses were most commonly enriched in both sexes. Negatively enriched gene sets found in both sexes included those associated with cell proliferation and cancer, estrogen responsiveness, and bile acid metabolism. Additionally, male mice had positive enrichment of gene sets associated with myogenesis, muscle development and intracellular calcium dynamics, whereas female mice had negative enrichment of gene sets associated with amino acid metabolism and collagen biosynthesis. Sucrose intake and sex interacted to positively enrich gene sets associated with adipogenesis and mitochondrial metabolism, and negatively enrich gene sets associated with myogenesis and breast development. In almost all cases, female sex had an additive effect on enrichment of gene sets that were enriched in both sexes.

For the subset of gene sets in the Hallmark database (Figure 4C and Supplementary File 4), sucrose intake led to enrichment of 65 distinct gene sets (45 positively enriched, 20 negatively enriched). In adipose tissue from male mice, sucrose enriched 16 Hallmark gene sets (13 positively enriched, three negatively enriched), whereas sucrose enriched 19 gene sets in adipose tissue from female mice (12 positively enriched, 7 negatively enriched). An additive effect of sex on sucrose intake was observed for 20 gene sets (16 positive, four negative), whereas ten gene sets showed sex:sucrose interactions (four positive, six negative). Positively-enriched pathways include those associated with fatty acid metabolism and adipogenesis, immunoregulation, mitochondrial bioenergetics and the cell cycle, whereas negatively-enriched pathways include those associated with estrogen signaling, bile acid metabolism and cell-junction organization and cell-cell communication.

### Sucrose Intake Alters The Transcriptional Regulation of Adrenergic Receptors in Female Mice

Since we were interested in identifying why male and female mice had divergent lipolysis adaptations in isolated adipocytes following sucrose intake, in addition to GSEA, we identified a subset of transcripts that encode either the enzymes and proteins that facilitate lipolysis (Supplementary Figure 3B and Supplementary File 2), or the receptors that regulate lipolysis activity either canonically (Supplementary Figure 3C and Supplementary File 2) or non-canonically (Supplementary Figure 3D and Supplementary File 2) (Braun et al., 2018). Transcripts encoding the enzymes and proteins that facilitate lipolysis were expressed similarly across all groups, although sucrose intake did increase the expression of transcripts encoding lipid droplet proteins known to moderate lipolytic activity (*Plin2* and *Plin5*). Expression of the transcript encoding the most abundant mouse adipose tissue β-AR, *Adrb3*, was reduced in female mice following sucrose intake (−11.7%, padj=0.013), as was the anti-lipolytic transcript *Adra2a* (−62.2%, padj=0.041). A sex:sucrose interaction effect was observed for the β-AR desensitizer *Arrb1* (padj=0.034), with female mice appearing to have reduced expression, although this did not attain statistical significance (−39.9%, padj=0.202). These findings identify reduced transcriptional regulation of adrenergic signaling as a potential mechanism responsible for the attenuated stimulated lipolysis response we observed in adipocytes from female mice after chronic sucrose intake.

### Sex Modulates the Effect of Chronic Sucrose Intake on The Microbiome

To gain insight into whether the increases in energy expenditure and altered substrate oxidation we observed following liquid sucrose intake were due to intrinsic changes to metabolism (Figure 1D and Supplemental Figure 1) or associated with alternations to the gut microbiome, we sequenced fecal samples from male and female mice from all four groups (Figure 5). At the phylum level, stool samples from female mice contained greater levels of Bacteroidetes than samples from male mice. At the family level, stool from females contained lower levels of unclassified *Rickettsiales* and *Clostridiales* and greater levels of *Lactobacillaceae*. Chronic sucrose intake increased the amount of *Lactobacillaceae* while decreasing *Prevotellaceae* (Figure 5A). Beta diversity was assessed by Bray Curtis and is shown as a principal coordinate analysis (PCoA; Figure 5B), where PCoA 1 explained 28% of diversity and discriminated sucrose vs controls. Dissimilar clustering was observed between groups (PERMANOVA R2 0.33, p=3.4^−4^; Anosim R = 0.512, p=1.0^−4^) while dispersion variability across groups was not significantly different (PERMDISP2 p=0.355). Redundancy analysis (RDA) identified groups as being significantly different (Variance = 117, F = 2.12, p=0.001) (Figure 5C). Alpha diversity was assessed by Shannon, Simpson’s and Evenness indexes (Figure 5D), where males showed greater diversity than females, and sucrose feeding decreased diversity in females, but not males. At the genus level, female mice that did not receive sucrose displayed greater *Lactobacillus*, but lower levels of unclassified *Clostridiales* compared to male mice that did not receive sucrose. Sucrose intake increased *Lactobacillus* and unclassified *Clostridiaceae* in females, while decreasing unclassified *Lachnospiraceae*, unclassified *Mycoplasmataceae*, *Prevotella*, *Candidatus Arthromitis* (Segmented filamentous bacteria), and AF12 compared to female mice that did not receive sucrose. (Figures 5 E-F). Predictive metagenomics showed multiple changes in metabolic pathways in female mice fed sucrose, while no differences were observed for male mice (Figure 5G). Specifically, amino acid metabolism was markedly decreased while fructose and mannose metabolism as well as glycolysis and gluconeogenesis KEGG pathways were upregulated in females following chronic sucrose intake. These differences were driven by significant changes in 52 microbial genes after FDR correction, while no significant differences were seen in males (Figures 5H-I, see also Supplemental File 5).

**Figure 5:**
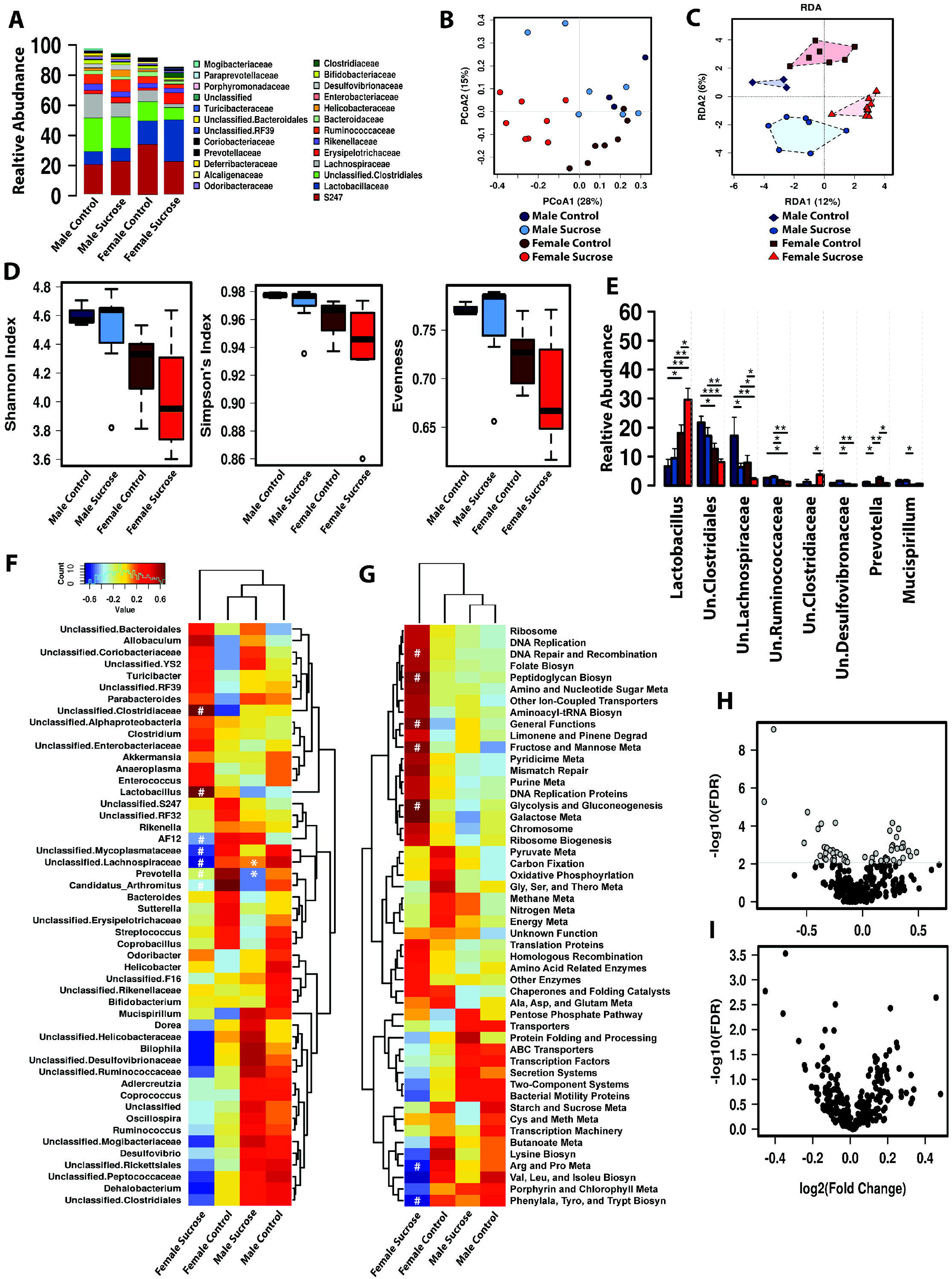
Chronic liquid sucrose intake has sex-specific effects on the microbiome. Microbiome relative abundance at the family level (A). Beta diversity by Bray Curtis displayed as principal coordinate analysis (PCoA) (B; PERMANOVA R2 0.33, p=3.4^−4^; Anosim R = 0.512, p=1.0^−4^; PERMDISP2 p= 0.355). Redundancy analysis (RDA) identified significantly dissimilar group clustering of beta diversity (C; Variance = 117, F = 2.12, p= 0.001). Alpha diversity assessed by Shannon, Simpson’s, and Evenness indexes (D). Significant changes in the top 20 most abundant genus by ANOVA (E). Heatmap of the 50 most abundant genus as calculated by spearman’s rank correlation coefficient (F; *p<0.05 vs male controls; ^#^p< 0.05 vs female controls). Heatmap of the 50 most abundant predicted metabolic pathways by PICRUSt displayed as calculated by spearman’s rank correlation coefficient (G; *p<0.05 vs male controls; ^#^p<0.05 vs female controls). Deseq2 volcano plot of KEGG genes in PICRUSt from female (H), and male (I) mice following sucrose intake compared to their respective control groups. All microbiome analyses n=3-8/group (see also: Supplemental File 5)

**Figure 6:**
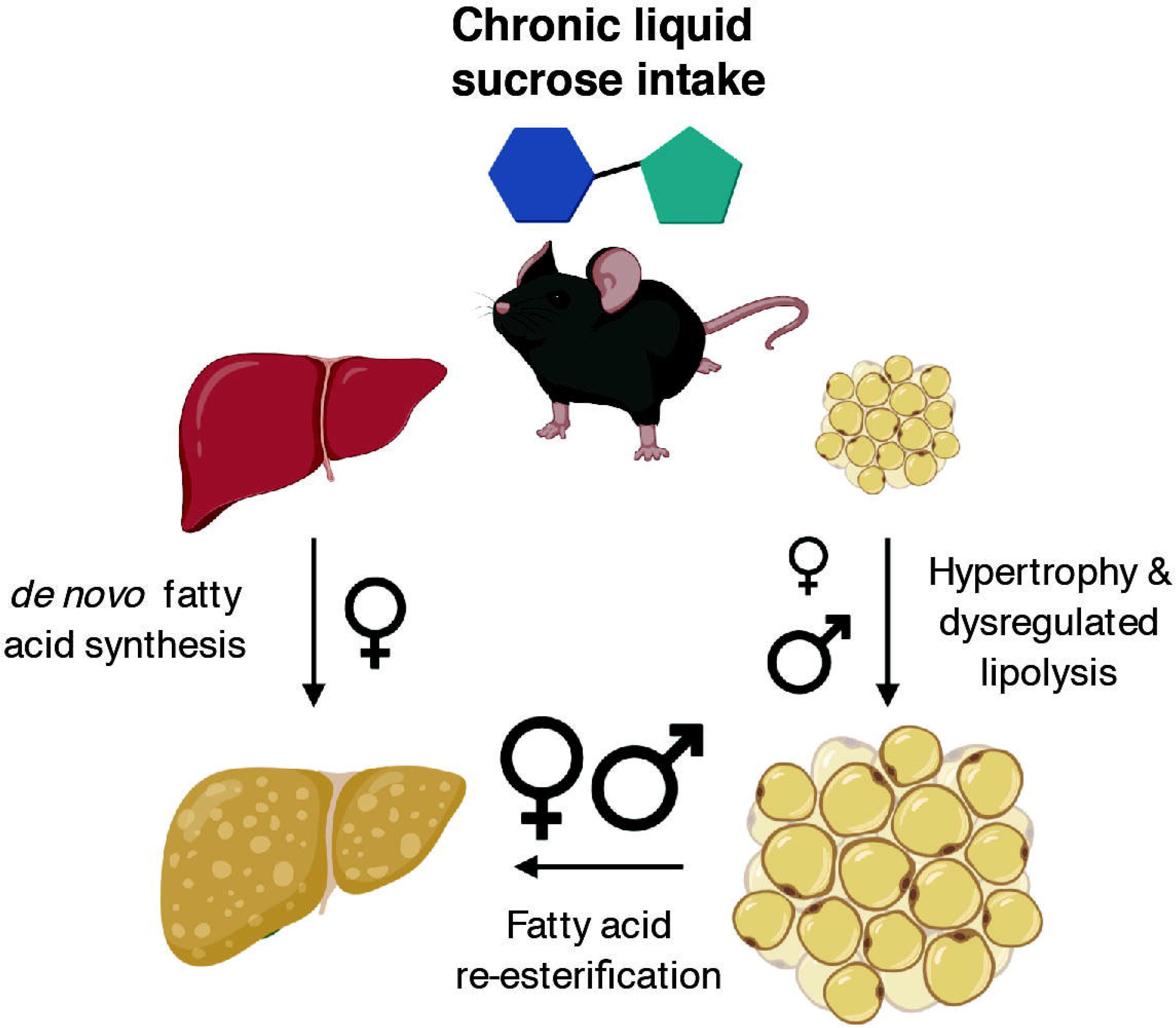
Summary of main findings. Chronic liquid sucrose intake at physiologically relevant concentrations has sexually dimorphic effects on liver and adipose tissue, differences that likely contribute to the severity of the resultant steatosis. In this study, we observed that male mice were able to convert excess dietary sucrose into fatty acids that were readily stored as triglyceride in the adipose tissue, whereas female mice were unable to expand their adipose tissue as effectively as males, leading to additional manufacturing (and storage) of new fatty acids in the liver. Our results suggest that liquid sucrose intake induces steatosis mainly via ‘spillover’ of fatty acids from dysregulated adipocyte lipolysis in both sexes, with additional contributions from *de novo* fatty acid synthesis occurring in females. Future studies will identify the mechanisms responsible for these divergent responses and allow us to better target therapeutic strategies for the treatment of NAFLD.

## Discussion

Here, we show that chronic intake of liquid sucrose at concentrations relevant to typical human consumption induces sexually dimorphic metabolic effects in liver, adipose tissue, and the microbiome; differences that likely contribute to the severity of sucrose-induced hepatic steatosis and the progression of NAFLD. We provide evidence that in the absence of systemic impairments in insulin responsiveness or glucose tolerance, sex is a moderating factor that interacts with the effects of high dietary sucrose intake to direct lipid storage and determine the contribution of *de novo* fatty acid synthesis to the hepatic triglyceride pool. These findings provide mechanistic insight into how sex influences the regulation of adipose-liver crosstalk in response to dietary sugar intake, and highlight the importance of extrahepatic metabolism in the pathogenesis of diet-induced steatosis and NAFLD.

Sucrose is a naturally occurring disaccharide comprised of equimolar amounts of the monosaccharides glucose and fructose. The refined form of sucrose, together with high-fructose corn syrup (which is not meaningfully different from sucrose in its composition), are the most commonly added sugars in the human diet (Coulston and Johnson, 2002; White, 2008). The ability of the fructose monosaccharide to bypass glycolytic regulation and proceed directly toward *de novo* fatty acid synthesis has long been speculated as the mechanism responsible for initiating NAFLD in response to the intake of dietary sugars (Mayes, 1993). Indeed, there are many reports in both preclinical (Crescenzo et al., 2013; Kawasaki et al., 2009; Nunes et al., 2014) and human populations (Le et al., 2009; Parks et al., 2008; Smajis et al., 2020) that support the assertion that fructose ingestion contributes to steatosis. However, fructose is rarely consumed in isolation, nor are added sugars consumed at the exaggerated concentrations frequently used to induce steatosis in animal models (Laughlin, 2014), and there is evidence in humans that suggests only those with existing metabolic disease are subject to fructose-induced gains in hepatic triglyceride when consumed in energy balance (Chung et al., 2014; Johnston et al., 2013; Le et al., 2006; Smajis et al., 2020). This lack of translatability, along with the long-supported notion that hepatic metabolism is tightly coupled to both endocrine signaling and systemic cues from metabolic intermediates generated endogenously by other tissues (Rui, 2014), makes it difficult to draw mechanistic conclusions about the initiating pathophysiology of NAFLD resulting from the intake of dietary sugars.

In the current study we have used a model of chronic sugar intake where chow-fed mice received sucrose in their drinking water at a concentration representative of many sugar sweetened beverages made for human consumption (Ventura et al., 2011). This ‘dose’ of sucrose was effective at increasing adiposity (Figure 1A-B) and inducing hepatic steatosis (Figure 2A) without causing systemic disruptions to insulin responsiveness or glucose tolerance (Supplementary Figure 2A-D). Notably, in contrast to previous studies (Mock et al., 2017; Schultz et al., 2015; Siddiqui et al., 2015), we observed that hepatic triglyceride accumulation following physiologically plausible sucrose intake was primarily driven by the reesterification of fatty acids derived from adipocyte lipolysis in male mice, and by a combination of reesterification and hepatic *de novo* fatty acid synthesis in female mice (Figures 2 and 3). Furthermore, we showed that ATGL-driven adipocyte lipolysis was required for sucrose-associated steatosis, although inhibition of lipolysis was only partially protective against steatosis in female mice (Figure 3D). Together with our observation that males had a higher capacity to store lipid in their adipose tissue than females (Figure 1A and B), these findings suggest that steatosis in the setting of high sugar intake may begin as a secondary consequence of dysregulated adipose tissue lipolysis and a reduced ability to store lipid in adipocytes, rather than a primary defect in hepatic metabolism *per se*. If this assertion is correct, it may help to explain why the most effective treatments for NAFLD to date have involved the reduction of adiposity, either via diet and exercise (Koutoukidis et al., 2019) or following surgical weight loss (Fakhry et al., 2019).

The preservation of systemic insulin responsiveness (and glucose tolerance; Supplementary Figure 2) in our model demonstrate a clear decoupling of the mechanisms responsible for steatosis and those responsible for impairments in insulin sensitivity following chronic sucrose intake and, in the case of female mice, overt obesity (Figure 1A). In humans, sugar sweetened beverage consumption is strongly associated with NAFLD (Arenaza et al., 2019; Asgari-Taee et al., 2019; Assy et al., 2008; Cahlin et al., 1973; Chen et al., 2019; Ma et al., 2015) but not always with obesity (Te Morenga et al., 2014). Additionally, NAFLD does not always present in parallel with obesity or insulin resistance (Shi et al., 2020) suggesting that the etiology of NAFLD likely deviates from that of other metabolic comorbidities. Our findings suggest that in the case of NAFLD resulting from high dietary sucrose intake, steatosis development is unlikely to be intrinsically linked with mechanisms responsible for systemic insulin resistance.

Our observation that male and female mice had mechanistically different responses to sucrose intake was not surprising. *In silico* modelling suggests NAFLD develops through distinct metabolic processes in males and females (Cvitanovic Tomas et al., 2018), whereas epidemiological data strongly support an increased prevalence of NAFLD in men (Ballestri et al., 2017; Group. et al., 2015), yet worsened NAFLD severity in women (Ballestri et al., 2017). Indeed, our observation that the severity of sucrose-induced steatosis was greatest in female mice (Figure 2A and B) is in line with previous studies that report worsened steatosis in female rats following high fructose (Hyer et al., 2019), or high fat, high fructose diets (Chukijrungroat et al., 2017). Together with the observation that women increase *de novo* fatty acid synthesis in response to acute fructose ingestion whereas men do not (Low et al., 2018), our findings add physiologic insight as to why females develop more severe steatosis than males in response to high dietary sugar intake.

In addition to highlighting the importance of metabolic crosstalk between liver and adipose tissue in the development of NAFLD, our study has identified a number of distinct transcriptional regulatory pathways that are induced in adipose tissue by chronic liquid sucrose intake (Figure 4 and Supplementary File 4).

Many of the pathways we identified were enriched in both male and female mice, whereas many gene sets also demonstrated an additive effect of sex. For example, we observed marked positive enrichment of gene sets associated with mitochondrial biogenesis and oxidative phosphorylation, an effect that was augmented in adipose tissue from female mice. Of the negatively enriched gene sets we observed, the downregulation of estrogen responsive gene sets in both male and female mice following chronic sucrose intake stood out as a potential mechanism that may, in part, explain some of the sexual dimorphism we observed.

Despite sucrose intake resulting in negative enrichment of estrogen signaling genes in adipose tissue from both sexes, a reduction in the transcript encoding Estrogen Receptor α (ERa; *Esr1*) was only observed in adipose tissue from female mice (Supplementary File 2). Estradiol interacting with ERα is a negative regulator of adipocyte hypertrophy (D’Eon et al., 2005; Davis et al., 2013). Thus, it is plausible that estrogen signaling becomes downregulated in adipose tissue as a means to increase adipocyte lipid storage in response to sucrose intake. In liver, estradiol has been shown to recruit ERα to the lipogenic genes *Fasn* and *Acaca*, resulting in their transcriptional inhibition (Qiu et al., 2017). Should this function translate across tissues, it could be speculated that lower endogenous concentrations of circulating estradiol in combination with attenuated basal expression of adipose tissue *Esr1* would assist male mice to rapidly increase their adiposity in response to sucrose intake, whereas the need to attenuate a higher basal *Esr1* expression would likely contribute to the milder adiposity gains we observe in female mice (Figure 1A and B).

Abrogation of circulating estradiol via ovariectomy (Chukijrungroat et al., 2017; Kamada et al., 2011), or selective loss of ERα (Della Torre et al., 2016; Qiu et al., 2017) in liver, each result in steatosis, with the latter showing sexual dimorphism in the steatosis response (Della Torre et al., 2016). Reductions in estrogen signaling in response to sucrose intake would be expected to increase hepatic *de novo* fatty acid synthesis via upregulation of *Fasn* and *Acaca* (Qiu et al., 2017). In support of this notion, liquid fructose ingestion (13% w/v) has been shown to cause a reduction in hepatic ERα protein expression concomitant with increases in *Fasn, Acaca*, and steatosis (Lin et al., 2019), whereas *de novo* fatty acid synthesis induced by sucrose, fructose or glucose has been shown to reduce the hepatocyte-produced sex hormone binding globulin (SHBG) (Selva et al., 2007). SHBG is thought to interact with ERα to regulate the transport of estrogens (and testosterone) into tissues (Hammond, 2016), and it’s concentrations are negatively associated with the severity of NAFLD and other metabolic disease risk-factors in humans (Kavanagh et al., 2013; Luo et al., 2018). In our model, hepatic expression of *Esr1* and *Shbg* are downregulated in female mice following sucrose intake but unchanged in males (not shown). Interestingly, SHBG has been found to increase lipolysis independently of sex steroids *in vitro* (Yamazaki et al., 2018), whereas its transcriptional coactivator HNF4a is regulated by the binding of fatty acids (Hertz et al., 1998), hinting at a possible role for SHBG in facilitating liver-adipose crosstalk in response to bioenergetic fluctuations. In the current study we observed similar *in vivo* lipolysis responses in both sexes (Figure 2A), but divergent *ex vivo* responses (Figure 3B), suggesting there may be a systemic signaling intermediate that helps facilitates the β-AR stimulated induction of lipolysis in adipocytes from female mice after chronic sucrose intake. Whether or not this intermediate is SHBG or something else entirely remains to be determined. Future studies will investigate how chronic sucrose intake triggers a reduction in adipose estrogen signaling, whether this effect is global, and if SHBG plays a role in facilitating crosstalk between the liver and adipose tissue in response to nutritional challenges.

In addition to the effects we observed on lipid metabolism, we (Figure 1D and Supplementary Figure 1) and others (Burke et al., 2018; Maekawa et al., 2017; Togo et al., 2019) have shown that energy expenditure increases in response to liquid sucrose intake. Given our observation of a disconnect between energy intake, energy expenditure and adiposity in the absence of any impairments to systemic insulin sensitivity following chronic liquid sucrose intake, we sought to determine if microbial metabolism might explain some of these metabolic adaptations. Consistent with this expectation, we observed that the microbiome of female mice that consumed sucrose was markedly altered in community diversity and membership, including increased *Lactobacillus* and *Clostridiaceae*. These changes correlated with alteration in predictive metabolic capacity, specifically amino acid and sugar metabolism pathways and genes. These data corroborate the sex specific differences in global response to sucrose metabolism and suggest the intestinal microbiome in female mice may be significant contributing factors to their distinct metabolic adaptation to sucrose intake.

### Limitations

It is important that we acknowledge some potential unintended consequences of our model of chronic sucrose intake. Mice receiving sucrose rapidly reduced their solid food intake (Figure 1C-D), presumably as a way of regulating their total caloric intake. In one respect, the lack of difference in energy intake between mice receiving sucrose and those in the control groups strengthens some of our observations because it prevented the confounding effect of excess energy intake. On the other hand, the macronutrient intake of the mice receiving sucrose was altered beyond the addition of sucrose. Based on the macronutrient breakdown provided by the manufacturer of the chow diet, male and female mice receiving sucrose consumed less protein (14.2% and 16.9%, respectively, compared to 25%) and less fat (9.6% and 11.5%, respectively, compared to 17%), and these differences may have contributed to some of the characteristics we observed in mice following chronic sucrose intake. For example, lower intake of protein and the subsequent reduction in branched chain amino acid availability may have contributed to the maintenance of systemic insulin sensitivity and glucose tolerance in mice receiving sucrose, as well as their lower lean mass (Mu et al., 2018; White et al., 2016). Nonetheless, mice still surpassed the lower threshold of fat and protein requirements needed for growth (1995; Goettsch, 1960) and many differences resulting from reduced intake of dietary fat or protein are likely to be overshadowed by the marked effects of sucrose in this model.

### Conclusion

We have shown that sex is a moderating factor in the regulation of lipid metabolism in both adipose tissue and liver. We demonstrate that mice consuming liquid sucrose at concentrations relevant to human consumption develop steatosis primarily as a result of dysregulated adipose tissue lipolysis and that the increased severity at which female mice develop steatosis is driven by upregulation of *de novo* fatty acid synthesis in addition to enhanced re-esterification. We speculate that the sexually dimorphic metabolic responses in adipose tissue and liver are likely due to impaired estrogen signaling either directly, through ERα, and/or indirectly, via SHBG or some other mechanism entirely. Future studies will investigate the mechanisms by which chronic sugar intake attenuates estrogen signaling and its role in liver-adipose crosstalk. In addition, our findings highlight the importance of preclinical studies that include both sexes in the experimental design, and the need for increased translatability in relation to preclinical research that involves the study of macronutrient metabolism. Lastly, when considering treatment strategies for reversing NAFLD, greater emphasis should be placed on extrahepatic lipid metabolism and the influence of sex and sex hormone signaling.

## Supporting information

Supplementary File 1: Primer sequences

Supplementary File 2: DEG

Supplementary File 3: StringDB

Supplementary File 4: GSEA

Supplementary File 5: Deseq2

## Acknowledgements

The authors would like to thank Daniel McCarty and the rest of the UTHSC Laboratory Animal Care Unit veterinary and husbandry team for their expert care of the mice that were studied for this report. We would also like to acknowledge the services contributed by Dr. Ilya Bederman from the Department of Pediatrics at Case Western Reserve University (determination of deuterium enrichment in blood samples), the UTHSC Molecular Research Center of Excellence (homogenization equipment and RNA bioanalysis) and the UTHSC Proteomics and Metabolomics Core Facility (quantification of total and deuterium-enriched lipid and glycerol). We would also like to thank all current and former members of the Han lab, and the UTHSC Metabolism Research Interest Group for insightful discussions that aided in the development of this work. Funding for this work was provided to EJS in the form of Junior Investigator Grants #65-1602 and # 64-1305 from Le Bonheur Children’s Hospital, the Children’s Foundation Research Institute and the Le Bonheur Associate Board. DB is the recipient of funding from the National Institute of Diabetes and Digestive and Kidney Diseases (RO1DK107535 and P30DK020572). MCM is a recipient of a Rackham Merit Fellowship. JCH is the recipient of funding from the Memphis Research Consortium and the Le Bonheur Children’s Foundation Research Institute.

## Data Availability

Microbiome data are available at DOI 10.17605/OSF.IO/F8A4R. RNAseq data will be available from the Gene Expression Omnibus upon manuscript publication.

## Author Contributions

Conceptualization: EJS, DB; Data curation: EJS, ASS, JFP, MAP; Formal analysis: EJS, JFP, MCM; Funding acquisition: EJS, DB, JCH; Investigation EJS, ASS, LM, CKG, AS, PKR, JFP; Project administration: EJS; Resources: EJS, JCH, MAP; Supervision: EJS, JCH; Validation: EJS, MAP, JFP; Visualization: EJS, JFP; Writing-original draft: EJS; Writing-review & editing: All authors; Submission approval: All authors

---

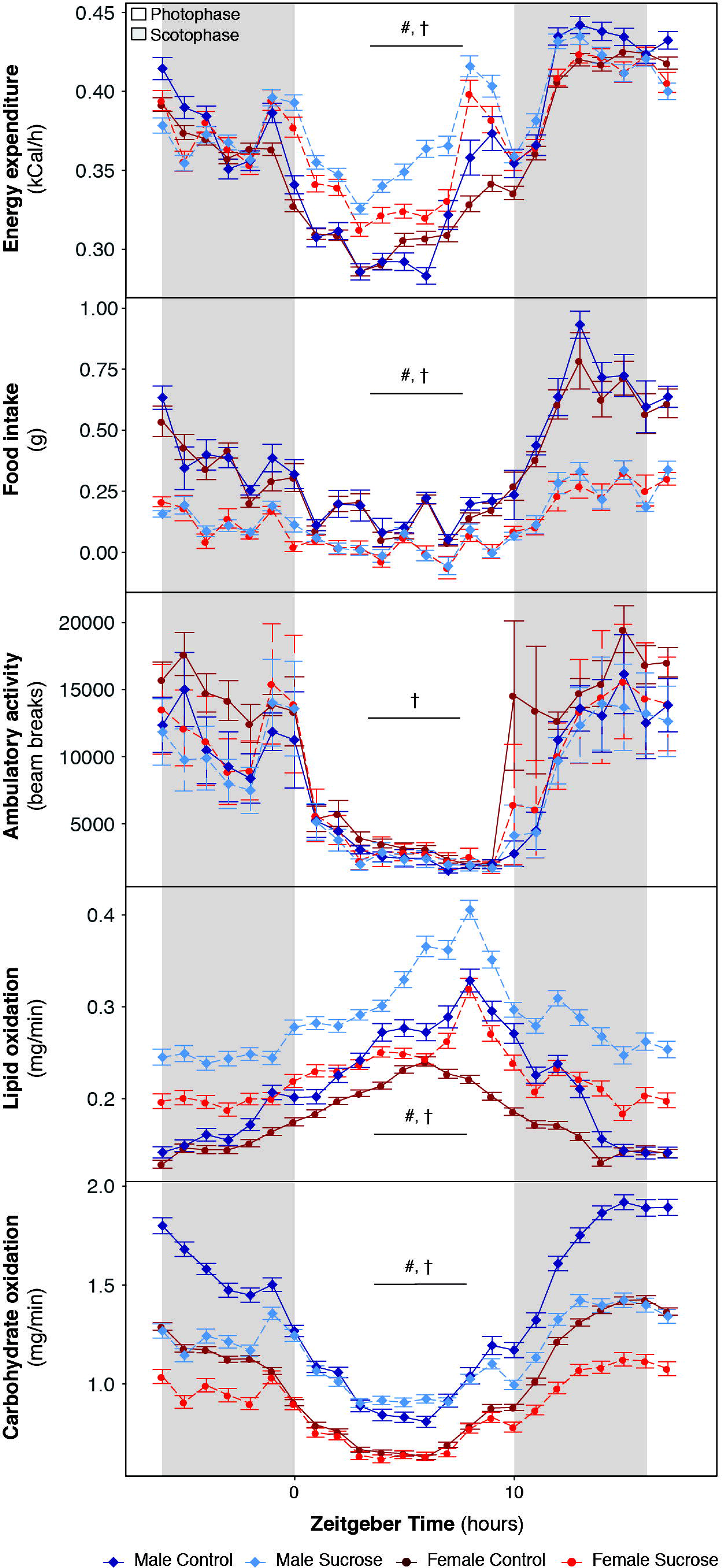

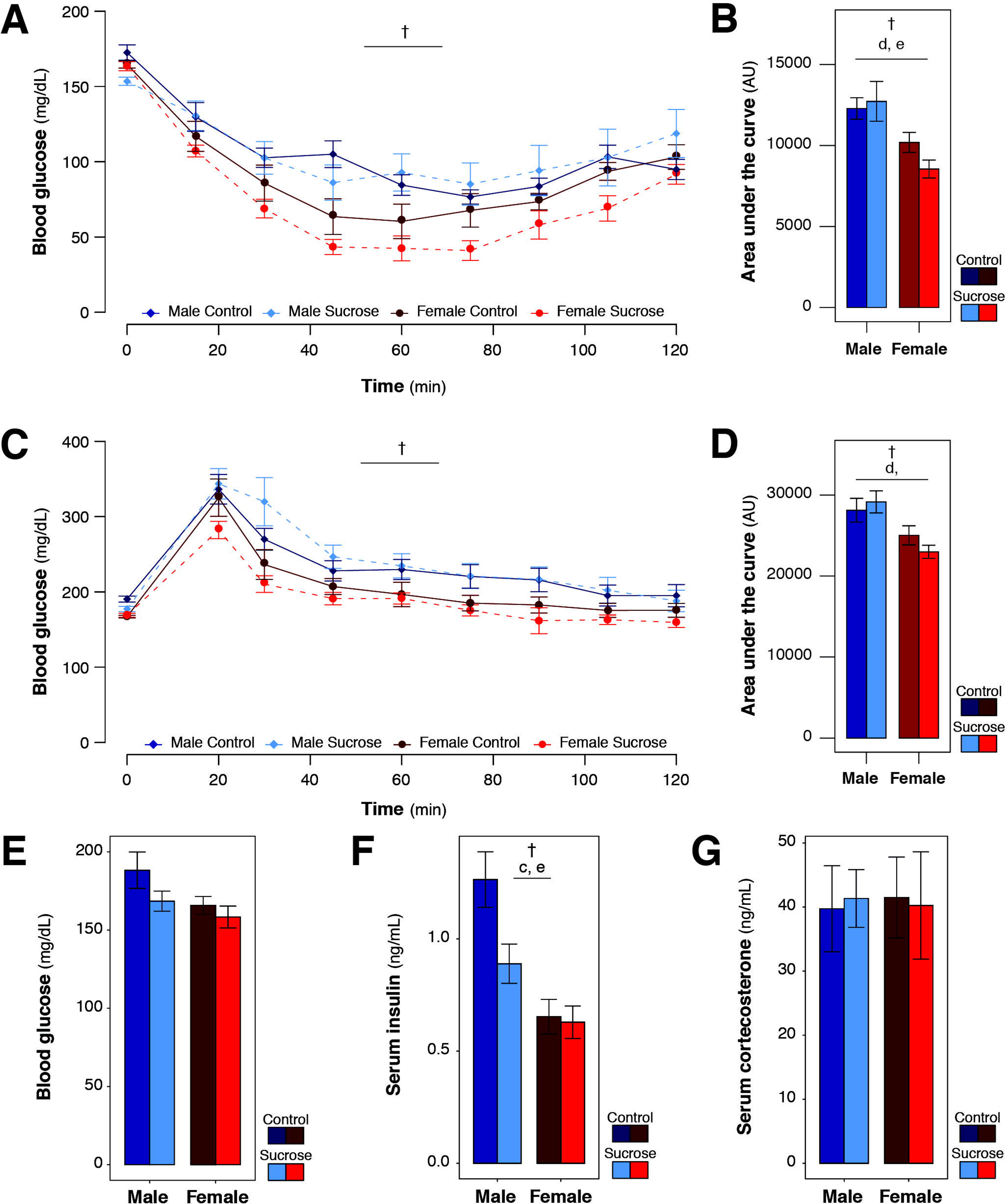

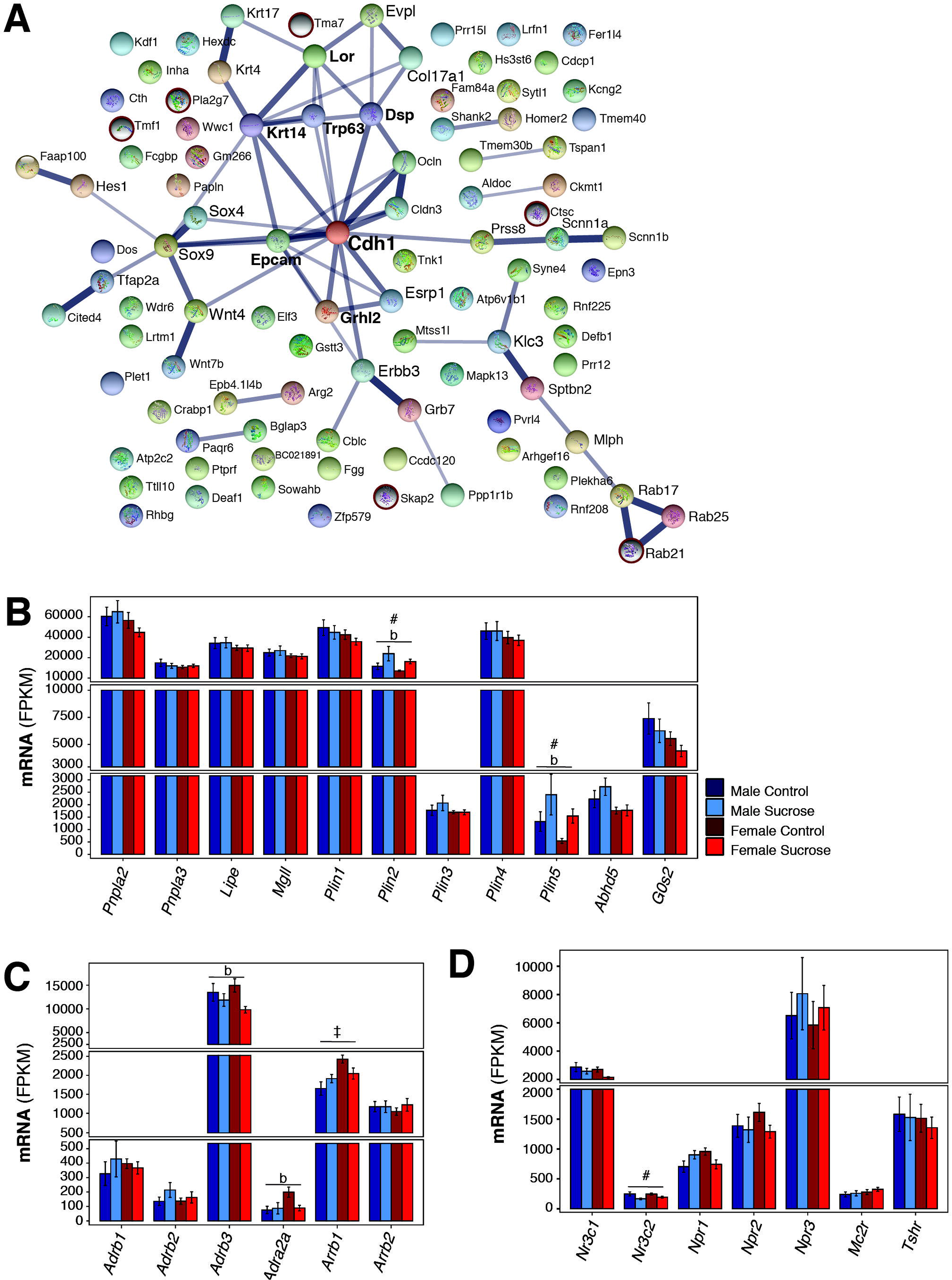

## References

Arenaza, L., Medrano, M., Oses, M., Huybrechts, I., Diez, I., Henriksson, H., and Labayen, I. (2019). Dietary determinants of hepatic fat content and insulin resistance in overweight/obese children: a cross-sectional analysis of the Prevention of Diabetes in Kids (PREDIKID) study. Br J Nutr 121, 1158–1165.

Armstrong, M.J., Adams, L.A., Canbay, A., and Syn, W.K. (2014). Extrahepatic complications of nonalcoholic fatty liver disease. Hepatology 59, 1174–1197.

Asgari-Taee, F., Zerafati-Shoae, N., Dehghani, M., Sadeghi, M., Baradaran, H.R., and Jazayeri, S. (2019). Association of sugar sweetened beverages consumption with non-alcoholic fatty liver disease: a systematic review and meta-analysis. Eur J Nutr 58, 1759–1769.

Assy, N., Nasser, G., Kamayse, I., Nseir, W., Beniashvili, Z., Djibre, A., and Grosovski, M. (2008). Soft drink consumption linked with fatty liver in the absence of traditional risk factors. Can J Gastroenterol 22, 811–816.

Bacon, B.R., Park, C.H., Fowell, E.M., and McLaren, C.E. (1984). Hepatic steatosis in rats fed diets with varying concentrations of sucrose. Fundam Appl Toxicol 4, 819–826.

Ballestri, S., Nascimbeni, F., Baldelli, E., Marrazzo, A., Romagnoli, D., and Lonardo, A. (2017). NAFLD as a Sexual Dimorphic Disease: Role of Gender and Reproductive Status in the Development and Progression of Nonalcoholic Fatty Liver Disease and Inherent Cardiovascular Risk. Adv Ther 34, 1291–1326.

Bates D, M.M., Bolker B, Walker S (2015). Fitting Linear Mixed-Effects Models Using lme4. Journal of Statistical Software 67, 1–48.

Bederman, I.R., Foy, S., Chandramouli, V., Alexander, J.C., and Previs, S.F. (2009). Triglyceride synthesis in epididymal adipose tissue: contribution of glucose and non-glucose carbon sources. J Biol Chem 284, 6101–6108.

Benjamini Y, H.Y. (1995). Controlling the false discovery rate: a practical and powerful approach to multiple testing. J R Stat Soc Ser B Methodol 57, 289–300.

Braun, K., Oeckl, J., Westermeier, J., Li, Y., and Klingenspor, M. (2018). Non-adrenergic control of lipolysis and thermogenesis in adipose tissues. J Exp Biol 221.

Brunengraber, D.Z., McCabe, B.J., Kasumov, T., Alexander, J.C., Chandramouli, V., and Previs, S.F. (2003). Influence of diet on the modeling of adipose tissue triglycerides during growth. Am J Physiol Endocrinol Metab 285, E917–925.

Bukowiecki, L.J., Lupien, J., Follea, N., and Jahjah, L. (1983). Effects of sucrose, caffeine, and cola beverages on obesity, cold resistance, and adipose tissue cellularity. Am J Physiol 244, R500–507.

Burke, S.J., Batdorf, H.M., Martin, T.M., Burk, D.H., Noland, R.C., Cooley, C.R., Karlstad, M.D., Johnson, W.D., and Collier, J.J. (2018). Liquid Sucrose Consumption Promotes Obesity and Impairs Glucose Tolerance Without Altering Circulating Insulin Levels. Obesity (Silver Spring) 26, 1188–1196.

Cahlin, E., Jonsson, J., Persson, B., Stakeberg, H., Bjorntorp, P., Gustafson, A., and Schersten, T. (1973). Sucrose feeding in man. Effects of substrate incorporation into hepatic triglycerides and phosphoglycerides in vitro and on removal of intravenous fat in patients with hyperlipoproteinemia. Scand J Clin Lab Invest 32, 21–33.

Caporaso, J.G., Kuczynski, J., Stombaugh, J., Bittinger, K., Bushman, F.D., Costello, E.K., Fierer, N., Pena, A.G., Goodrich, J.K., Gordon, J.I., et al. (2010). QIIME allows analysis of high-throughput community sequencing data. Nat Methods 7, 335–336.

Chen, G.C., Huang, C.Y., Chang, M.Y., Chen, C.H., Chen, S.W., Huang, C.J., and Chao, P.M. (2011). Two unhealthy dietary habits featuring a high fat content and a sucrose-containing beverage intake, alone or in combination, on inducing metabolic syndrome in Wistar rats and C57BL/6J mice. Metabolism 60, 155–164.

Chen, H., Wang, J., Li, Z., Lam, C.W.K., Xiao, Y., Wu, Q., and Zhang, W. (2019). Consumption of Sugar-Sweetened Beverages Has a Dose-Dependent Effect on the Risk of Non-Alcoholic Fatty Liver Disease: An Updated Systematic Review and Dose-Response Meta-Analysis. Int J Environ Res Public Health 16.

Chong, M.F., Fielding, B.A., and Frayn, K.N. (2007). Mechanisms for the acute effect of fructose on postprandial lipemia. Am J Clin Nutr 85, 1511–1520.

Choo, V.L., and Sievenpiper, J.L. (2015). The ecologic validity of fructose feeding trials: supraphysiological feeding of fructose in human trials requires careful consideration when drawing conclusions on cardiometabolic risk. Front Nutr 2, 12.

Chukijrungroat, N., Khamphaya, T., Weerachayaphorn, J., Songserm, T., and Saengsirisuwan, V. (2017). Hepatic FGF21 mediates sex differences in high-fat high-fructose diet-induced fatty liver. Am J Physiol Endocrinol Metab 313, E203–E212.

Chung, M., Ma, J., Patel, K., Berger, S., Lau, J., and Lichtenstein, A.H. (2014). Fructose, high-fructose corn syrup, sucrose, and nonalcoholic fatty liver disease or indexes of liver health: a systematic review and meta-analysis. Am J Clin Nutr 100, 833–849.

Coulston, A.M., and Johnson, R.K. (2002). Sugar and sugars: myths and realities. J Am Diet Assoc 102, 351–353.

Crescenzo, R., Bianco, F., Falcone, I., Coppola, P., Liverini, G., and Iossa, S. (2013). Increased hepatic de novo lipogenesis and mitochondrial efficiency in a model of obesity induced by diets rich in fructose. Eur J Nutr 52, 537–545.

Cvitanovic Tomas, T., Urlep, Z., Moskon, M., Mraz, M., and Rozman, D. (2018). LiverSex Computational Model: Sexual Aspects in Hepatic Metabolism and Abnormalities. Front Physiol 9, 360.

D’Eon, T.M., Souza, S.C., Aronovitz, M., Obin, M.S., Fried, S.K., and Greenberg, A.S. (2005). Estrogen regulation of adiposity and fuel partitioning. Evidence of genomic and non-genomic regulation of lipogenic and oxidative pathways. J Biol Chem 280, 35983–35991.

Davis, K.E., Neinast, D.N., Sun, K., Skiles, W.M., Bills, J.D., Zehr, J.A., Zeve, D., Hahner, L.D., Cox, D.W., Gent, L.M., et al. (2013). The sexually dimorphic role of adipose and adipocyte estrogen receptors in modulating adipose tissue expansion, inflammation, and fibrosis. Mol Metab 2, 227–242.

Della Torre, S., Mitro, N., Fontana, R., Gomaraschi, M., Favari, E., Recordati, C., Lolli, F., Quagliarini, F., Meda, C., Ohlsson, C., et al. (2016). An Essential Role for Liver ERalpha in Coupling Hepatic Metabolism to the Reproductive Cycle. Cell Rep 15, 360–371.

Diraison, F., Pachiaudi, C., and Beylot, M. (1997). Measuring lipogenesis and cholesterol synthesis in humans with deuterated water: use of simple gas chromatographic/mass spectrometric techniques. J Mass Spectrom 32, 81–86.

Eguchi, J., Wang, X., Yu, S., Kershaw, E.E., Chiu, P.C., Dushay, J., Estall, J.L., Klein, U., Maratos-Flier, E., and Rosen, E.D. (2011). Transcriptional control of adipose lipid handling by IRF4. Cell Metab 13, 249–259.

Fakhry, T.K., Mhaskar, R., Schwitalla, T., Muradova, E., Gonzalvo, J.P., and Murr, M.M. (2019). Bariatric surgery improves nonalcoholic fatty liver disease: a contemporary systematic review and metaanalysis. Surg Obes Relat Dis 15, 502–511.

Fox J, W.S. (2019). An R Companion to Applied Regression, Third edition.. (Thousand Oaks CA: Sage,).

Goettsch, M. (1960). Comparative Protein Requirement of the Rat and Mouse for Growth, Reproduction and Lactation Using Casein Diets. The Journal of Nutrition 70, 307–312.

Non-alcoholic Fatty Liver Disease Study Group, Lonardo, A., Bellentani, S., Argo, C.K., Ballestri, S., Byrne, C.D., Caldwell, S.H., Cortez-Pinto, H., Grieco, A., Machado, M.V., et al. (2015). Epidemiological modifiers of non-alcoholic fatty liver disease: Focus on high-risk groups. Dig Liver Dis 47, 997–1006.

Hammond, G.L. (2016). Plasma steroid-binding proteins: primary gatekeepers of steroid hormone action. J Endocrinol 230, R13–25.

Hellerstein, M.K. (1999). De novo lipogenesis in humans: metabolic and regulatory aspects. Eur J Clin Nutr 53 Suppl 1, S53–65.

Hertz, R., Magenheim, J., Berman, I., and Bar-Tana, J. (1998). Fatty acyl-CoA thioesters are ligands of hepatic nuclear factor-4alpha. Nature 392, 512–516.

Hughes, J.B., Hellmann, J. J., Ricketts, T. H., and Bohannan, B. J. (2001). Counting the uncountable: statistical approaches to estimating microbial diversity. Appl. Environ. Microbiol. 67, 4399–4406.

Hyer, M.M., Dyer, S.K., Kloster, A., Adrees, A., Taetzsch, T., Feaster, J., Valdez, G., and Neigh, G.N. (2019). Sex modifies the consequences of extended fructose consumption on liver health, motor function, and physiological damage in rats. Am J Physiol Regul Integr Comp Physiol 317, R903–R911.

Johnston, R.D., Stephenson, M.C., Crossland, H., Cordon, S.M., Palcidi, E., Cox, E.F., Taylor, M.A., Aithal, G.P., and Macdonald, I.A. (2013). No difference between high-fructose and high-glucose diets on liver triacylglycerol or biochemistry in healthy overweight men. Gastroenterology 145, 1016–1025 e1012.

Kamada, Y., Kiso, S., Yoshida, Y., Chatani, N., Kizu, T., Hamano, M., Tsubakio, M., Takemura, T., Ezaki, H., Hayashi, N., et al. (2011). Estrogen deficiency worsens steatohepatitis in mice fed high-fat and high-cholesterol diet. Am J Physiol Gastrointest Liver Physiol 301, G1031–1043.

Kavanagh, K., Espeland, M.A., Sutton-Tyrrell, K., Barinas-Mitchell, E., El Khoudary, S.R., and Wildman, R.P. (2013). Liver fat and SHBG affect insulin resistance in midlife women: the Study of Women’s Health Across the Nation (SWAN). Obesity (Silver Spring) 21, 1031–1038.

Kawasaki, T., Igarashi, K., Koeda, T., Sugimoto, K., Nakagawa, K., Hayashi, S., Yamaji, R., Inui, H., Fukusato, T., and Yamanouchi, T. (2009). Rats fed fructose-enriched diets have characteristics of nonalcoholic hepatic steatosis. J Nutr 139, 2067–2071.

Kim, D., Kim, W.R., Kim, H.J., and Therneau, T.M. (2013). Association between noninvasive fibrosis markers and mortality among adults with nonalcoholic fatty liver disease in the United States. Hepatology 57, 1357–1365.

Koutoukidis, D.A., Astbury, N.M., Tudor, K.E., Morris, E., Henry, J.A., Noreik, M., Jebb, S.A., and Aveyard, P. (2019). Association of Weight Loss Interventions With Changes in Biomarkers of Nonalcoholic Fatty Liver Disease: A Systematic Review and Meta-analysis. JAMA Intern Med.

Laughlin, M.R. (2014). Normal roles for dietary fructose in carbohydrate metabolism. Nutrients 6, 3117–3129.

Le, K.A., Faeh, D., Stettler, R., Ith, M., Kreis, R., Vermathen, P., Boesch, C., Ravussin, E., and Tappy, L. (2006). A 4-wk high-fructose diet alters lipid metabolism without affecting insulin sensitivity or ectopic lipids in healthy humans. Am J Clin Nutr 84, 1374–1379.

Le, K.A., Ith, M., Kreis, R., Faeh, D., Bortolotti, M., Tran, C., Boesch, C., and Tappy, L. (2009). Fructose overconsumption causes dyslipidemia and ectopic lipid deposition in healthy subjects with and without a family history of type 2 diabetes. Am J Clin Nutr 89, 1760–1765.

Lin, R., Jia, Y., Wu, F., Meng, Y., Sun, Q., and Jia, L. (2019). Combined Exposure to Fructose and Bisphenol A Exacerbates Abnormal Lipid Metabolism in Liver of Developmental Male Rats. Int J Environ Res Public Health 16.

Lonardo, A., Nascimbeni, F., Ballestri, S., Fairweather, D., Win, S., Than, T.A., Abdelmalek, M.F., and Suzuki, A. (2019). Sex Differences in Nonalcoholic Fatty Liver Disease: State of the Art and Identification of Research Gaps. Hepatology 70, 1457–1469.

Love, M.I., Huber, W., Anders, S. (2014). Moderated estimation of fold change and dispersion for RNA-seq data with DESeq2. Genome Biology 15.

Low, W.S., Cornfield, T., Charlton, C.A., Tomlinson, J.W., and Hodson, L. (2018). Sex Differences in Hepatic De Novo Lipogenesis with Acute Fructose Feeding. Nutrients 10.

Lozupone, C., Lladser, M.E., Knights, D., Stombaugh, J., and Knight, R. (2011). UniFrac: an effective distance metric for microbial community comparison. ISME J 5, 169–172.

Luo, J., Chen, Q., Shen, T., Wang, X., Fang, W., Wu, X., Yuan, Z., Chen, G., Ling, W., and Chen, Y. (2018). Association of sex hormone-binding globulin with nonalcoholic fatty liver disease in Chinese adults. Nutr Metab (Lond) 15, 79.

Lusk, G. (1924). Animal Calorimetry: Twenty-Fourth Paper. Analysis of the Oxidation of Mixtures of Carbohydrate and Fat. Journal of Biologial Chemistry 59, 41–42.

Ma, J., Fox, C.S., Jacques, P.F., Speliotes, E.K., Hoffmann, U., Smith, C.E., Saltzman, E., and McKeown, N.M. (2015). Sugar-sweetened beverage, diet soda, and fatty liver disease in the Framingham Heart Study cohorts. J Hepatol 63, 462–469.

Maekawa, R., Seino, Y., Ogata, H., Murase, M., Iida, A., Hosokawa, K., Joo, E., Harada, N., Tsunekawa, S., Hamada, Y., et al. (2017). Chronic high-sucrose diet increases fibroblast growth factor 21 production and energy expenditure in mice. J Nutr Biochem 49, 71–79.

Mangiafico, S.S. (2016). Summary and Analysis of Extension Program Evaluation in R.

Mayes, P.A. (1993). Intermediary metabolism of fructose. Am J Clin Nutr 58, 754S–765S.

Mock, K., Lateef, S., Benedito, V.A., and Tou, J.C. (2017). High-fructose corn syrup-55 consumption alters hepatic lipid metabolism and promotes triglyceride accumulation. J Nutr Biochem 39, 32–39.

Mu, W.C., VanHoosier, E., Elks, C.M., and Grant, R.W. (2018). Long-Term Effects of Dietary Protein and Branched-Chain Amino Acids on Metabolism and Inflammation in Mice. Nutrients 10.

National Research Council (US) Subcommittee on Laboratory Animal Nutrition (1995). In Nutrient Requirements of Laboratory Animals: Fourth Revised Edition, 1995 (Washington (DC)).

Nunes, P.M., Wright, A.J., Veltien, A., van Asten, J.J., Tack, C.J., Jones, J.G., and Heerschap, A. (2014). Dietary lipids do not contribute to the higher hepatic triglyceride levels of fructose-compared to glucose-fed mice. FASEB J 28, 1988–1997.

Oliveira, L.S., Santos, D.A., Barbosa-da-Silva, S., Mandarim-de-Lacerda, C.A., and Aguila, M.B. (2014). The inflammatory profile and liver damage of a sucrose-rich diet in mice. J Nutr Biochem 25, 193–200.

Parks, E.J., Skokan, L.E., Timlin, M.T., and Dingfelder, C.S. (2008). Dietary sugars stimulate fatty acid synthesis in adults. J Nutr 138, 1039–1046.

Peronnet, F., and Massicotte, D. (1991). Table of nonprotein respiratory quotient: an update. Can J Sport Sci 16, 23–29.

Qiu, S., Vazquez, J.T., Boulger, E., Liu, H., Xue, P., Hussain, M.A., and Wolfe, A. (2017). Hepatic estrogen receptor alpha is critical for regulation of gluconeogenesis and lipid metabolism in males. Sci Rep 7, 1661.

Ritze, Y., Bardos, G., D’Haese, J.G., Ernst, B., Thurnheer, M., Schultes, B., and Bischoff, S.C. (2014). Effect of high sugar intake on glucose transporter and weight regulating hormones in mice and humans. PLoS One 9, e101702.

Rui, L. (2014). Energy metabolism in the liver. Compr Physiol 4, 177–197.

Sanyal, A.J. (2019). Past, present and future perspectives in nonalcoholic fatty liver disease. Nat Rev Gastroenterol Hepatol 16, 377–386.

Schultz, A., Barbosa-da-Silva, S., Aguila, M.B., and Mandarim-de-Lacerda, C.A. (2015). Differences and similarities in hepatic lipogenesis, gluconeogenesis and oxidative imbalance in mice fed diets rich in fructose or sucrose. Food Funct 6, 1684–1691.

Selva, D.M., Hogeveen, K.N., Innis, S.M., and Hammond, G.L. (2007). Monosaccharide-induced lipogenesis regulates the human hepatic sex hormone-binding globulin gene. J Clin Invest 117, 3979–3987.

Sergushichev, A (2016). An algorithm for fast preranked gene set enrichment analysis using cumulative statistic calculation. bioRxiv. doi: https://doi.org/10.1101/060012

Shi, Y., Wang, Q., Sun, Y., Zhao, X., Kong, Y., Ou, X., Jia, J., Wu, S., and You, H. (2020). The Prevalence of Lean/Nonobese Nonalcoholic Fatty Liver Disease: A Systematic Review and MetaAnalysis. J Clin Gastroenterol 54, 378–387.

Siddiqui, R.A., Xu, Z., Harvey, K.A., Pavlina, T.M., Becker, M.J., and Zaloga, G.P. (2015). Comparative study of the modulation of fructose/sucrose-induced hepatic steatosis by mixed lipid formulations varying in unsaturated fatty acid content. Nutr Metab (Lond) 12, 41.

Sitnick, M.T., Basantani, M.K., Cai, L., Schoiswohl, G., Yazbeck, C.F., Distefano, G., Ritov, V., DeLany, J.P., Schreiber, R., Stolz, D.B., et al. (2013). Skeletal muscle triacylglycerol hydrolysis does not influence metabolic complications of obesity. Diabetes 62, 3350–3361.

Smajis, S., Gajdosik, M., Pfleger, L., Traussnigg, S., Kienbacher, C., Halilbasic, E., Ranzenberger-Haider, T., Stangl, A., Beiglbock, H., Wolf, P., et al. (2020). Metabolic effects of a prolonged, very-high-dose dietary fructose challenge in healthy subjects. Am J Clin Nutr 111, 369–377.

Smith, U., Cahlin, E., and Schersten, T. (1973). Sucrose feeding in man. Effects on lipolysis and antilipolytic action of insulin in the adipose tissue. Acta Med Scand 194, 147–150.

Softic, S., Meyer, J.G., Wang, G.X., Gupta, M.K., Batista, T.M., Lauritzen, H., Fujisaka, S., Serra, D., Herrero, L., Willoughby, J., et al. (2019). Dietary Sugars Alter Hepatic Fatty Acid Oxidation via Transcriptional and Post-translational Modifications of Mitochondrial Proteins. Cell Metab 30, 735–753 e734.

Soria, A., D’Alessandro, M.E., and Lombardo, Y.B. (2001). Duration of feeding on a sucrose-rich diet determines metabolic and morphological changes in rat adipocytes. J Appl Physiol (1985) 91, 2109–2116.

Spruss, A., and Bergheim, I. (2009). Dietary fructose and intestinal barrier: potential risk factor in the pathogenesis of nonalcoholic fatty liver disease. J Nutr Biochem 20, 657–662.

Sun, S.Z., and Empie, M.W. (2012). Fructose metabolism in humans – what isotopic tracer studies tell us. Nutr Metab (Lond) 9, 89.

Szklarczyk, D., Gable, A.L., Lyon, D., Junge, A., Wyder, S., Huerta-Cepas, J., Simonovic, M., Doncheva, N.T., Morris, J.H., Bork, P., et al. (2019). STRING v11: protein-protein association networks with increased coverage, supporting functional discovery in genome-wide experimental datasets. Nucleic Acids Res 47, D607–D613.

Te Morenga, L.A., Howatson, A.J., Jones, R.M., and Mann, J. (2014). Dietary sugars and cardiometabolic risk: systematic review and meta-analyses of randomized controlled trials of the effects on blood pressure and lipids. Am J Clin Nutr 100, 65–79.

Togo, J., Hu, S., Li, M., Niu, C., and Speakman, J.R. (2019). Impact of dietary sucrose on adiposity and glucose homeostasis in C57BL/6J mice depends on mode of ingestion: liquid or solid. Mol Metab 27, 22–32.

van Hall, G., Sacchetti, M., Radegran, G., and Saltin, B. (2002). Human skeletal muscle fatty acid and glycerol metabolism during rest, exercise and recovery. J Physiol 543, 1047–1058.

Ventura, E.E., Davis, J.N., and Goran, M.I. (2011). Sugar content of popular sweetened beverages based on objective laboratory analysis: focus on fructose content. Obesity (Silver Spring) 19, 868–874.

White, J.S. (2008). Straight talk about high-fructose corn syrup: what it is and what it ain’t. Am J Clin Nutr 88, 1716S–1721S.

White, P.J., Lapworth, A.L., An, J., Wang, L., McGarrah, R.W., Stevens, R.D., Ilkayeva, O., George, T., Muehlbauer, M.J., Bain, J.R., et al. (2016). Branched-chain amino acid restriction in Zucker-fatty rats improves muscle insulin sensitivity by enhancing efficiency of fatty acid oxidation and acylglycine export. Mol Metab 5, 538–551.

Yamazaki, H., Kushiyama, A., Sakoda, H., Fujishiro, M., Yamamotoya, T., Nakatsu, Y., Kikuchi, T., Kaneko, S., Tanaka, H., and Asano, T. (2018). Protective Effect of Sex Hormone-Binding Globulin against Metabolic Syndrome: In Vitro Evidence Showing Anti-Inflammatory and Lipolytic Effects on Adipocytes and Macrophages. Mediators Inflamm 2018, 3062319.

Younossi, Z.M., Koenig, A.B., Abdelatif, D., Fazel, Y., Henry, L., and Wymer, M. (2016). Global epidemiology of nonalcoholic fatty liver disease-Meta-analytic assessment of prevalence, incidence, and outcomes. Hepatology 64, 73–84.

Zakrzewski, M., Proietti, C., Ellis, J. J., Hasan, S., Brion, M.-J., Berger, B., and Krause, L. (2017). Calypso: a user-friendly web-server for mining and visualizing microbiome-environment interactions. Bioinformatics 33, 782–783.

